# No evidence of immunosurveillance in mutation-hotspot driven clonal haematopoiesis

**DOI:** 10.1101/2024.09.27.615394

**Authors:** Barbara Walkowiak, Hamish AJ MacGregor, Jamie R Blundell

## Abstract

The theory of immunosurveillance posits that T-cells can selectively eliminate clones harbouring non-self antigens generated by somatic mutations. There is considerable evidence supporting the role of immune surveillance in cancer. Whether immunosurveillance imposes a negative selective pressure on pre-cancerous clones, however, is not well established. Here, we studied the association between MHC-variant binding and risk of clonal haematopoiesis (CH), a pre-cancer state in the blood driven by expansions of mutant haematopoietic stem cells (HSCs). We predicted MHC binding affinity towards 40 known CH hotspot variants in 380,000 UK Biobank participants, and examined the relationship between predicted binding to each variant and risk of its expansion in the blood. Despite being well powered to detect subtle differences in selective pressure, we did not find associations between predicted MHC binding and CH prevalence for any of the hotspot variants. In individuals in whom we identified CH, there was no relationship between predicted binding affinity to the variant and size of the clone. Overall, we do not find evidence for the MHC genotype to be a factor that affects which somatic variants expand in CH, suggesting a limited role for immunosurveillance in shaping the genetic diversity of the blood.

## INTRODUCTION

According to the theory of immunosurveillance, the immune system is able to identify and eliminate cells carrying somatic mutations, thus preventing potentially cancerous clones from expanding ^1^. All cells display peptides generated from their proteome on the surface, allowing the immune system to detect cells that present non-self peptides. Peptide presentation requires its binding to the class I major histocompatibility complex (MHC-I). The resulting peptide-MHC (p-MHC) complex is trafficked to the cell surface, where it can be recognised by CD8+ T cells ^2^. Whether a given peptide is presented on the surface is influenced by its binding affinity to the MHC, with strongly binding peptides being more likely to form the p-MHC complex ^3^. MHC alleles are highly polymorphic in the human population ^4^, therefore a peptide may be differentially presented between individuals who carry different MHC alleles. This has been suggested to influence the expansion of driver mutations in cancer. Cancers reportedly carry driver mutations that are anomalously poor binders to the patients MHC ^5^ and particular MHC alleles predicted to bind mutant peptides with high affinity are under-represented in some cancers ^6^. In addition to MHC I, recognition of neoantigens presented on the MHC II complex, which is mainly expressed in hematopoietic and immune cells ^7,8^, by CD4+ T cells can contribute to detection and elimination of cancer clones ^9,10^.

While extensively studied in the context of cancer ^7,11–15^, it is not fully understood when immunosurveillance of somatic clones begins and to what extent it shapes somatic evolution in ageing, pre-cancerous tissues. Incidence of cancer increases in immuno-compromised individuals ^16^, suggesting that the immune system can prevent early clonal expansions from transforming into clinically observable disease. While no signatures of negative selection or immune system activity were identified in healthy somatic tissues which bear a significant burden of mutations ^17,18^, tissues in early pre-cancer stages show signs of immune infiltration ^19^. This could be related to the observation that immune system engagement may be dependent on reaching a minimum fraction of cells which bear the neo-antigen in the tissue ^20^. However, studying early immune surveillance has been challenging due to small sample sizes and focus on small clonal expansions. Therefore, datasets where a large number of clonal expansions can be measured quantitatively are needed to comprehensively address the problem.

We reasoned that the role of immune surveillance during somatic evolution could be investigated by assessing whether the evolutionary trajectory of a variant is influenced by the capacity of the immune system to recognise it. We examined this in clonal haematopoiesis (CH), a pre-cancer state which results from expansion of haematopoietic stem cells (HSCs) and is associated with an increased risk of haematological malignancies ^21,22^. CH can be identified in blood exome sequencing data, available for cohorts such as the UK Biobank (UKB), enabling the comparison of expansions of a specific driver hotspot mutation across a large number of individuals who differ in their capacity to present this mutation, providing us with considerable power to detect potential associations. In particular, in individuals whose MHC presents a given variant, we may expect expansions of clones carrying this variant to be more limited. In contrast, in individuals whose MHC does not bind peptides with this variant, there will be no immune recognition, and so expansions will not be restricted. At the population level, individuals who present a given mutation well are therefore predicted to have an overall lower prevalence of clones harbouring this variant.

## RESULTS

### Association between MHC binding and CH prevalence

The UK Biobank contains whole exome sequencing data from around 450,000 people between the ages of 40 and 70. Due to the depth of sequencing (median depth of 50x) and known propensity for CH to occur at certain positions in the genome (CH “hotspots”), somatic clones can be robustly identified in these data, allowing us to explore the potential relationship between MHC presentation and CH at considerable scale. If immunosurveillence were able to exert negative selective pressure on somatic clones, we reasoned that individuals predicted to have stronger MHC binding to a specific variant peptide may be at lower risk of developing CH driven by that variant (Fig. 1A). To test this, we obtained imputed MHC genotypes from cancer-free individuals in the UKB ^23^ (MHC class I: n=384, 600; class II: n=327, 961) and used *NetMHCpan-4*.*1* ^24^ to make computational predictions of the MHC I binding affinity between each individual’s MHC I alleles and peptides carrying 40 recurrent missense and truncating variants in 11 common CH driver genes ^25^. Each person-variant pair was assigned a single MHC binding score defined as the strongest-binding peptide that covers the variant measured via elution percentage rank score across all MHC alleles present in that individual (see methods).

**Fig. 1.**
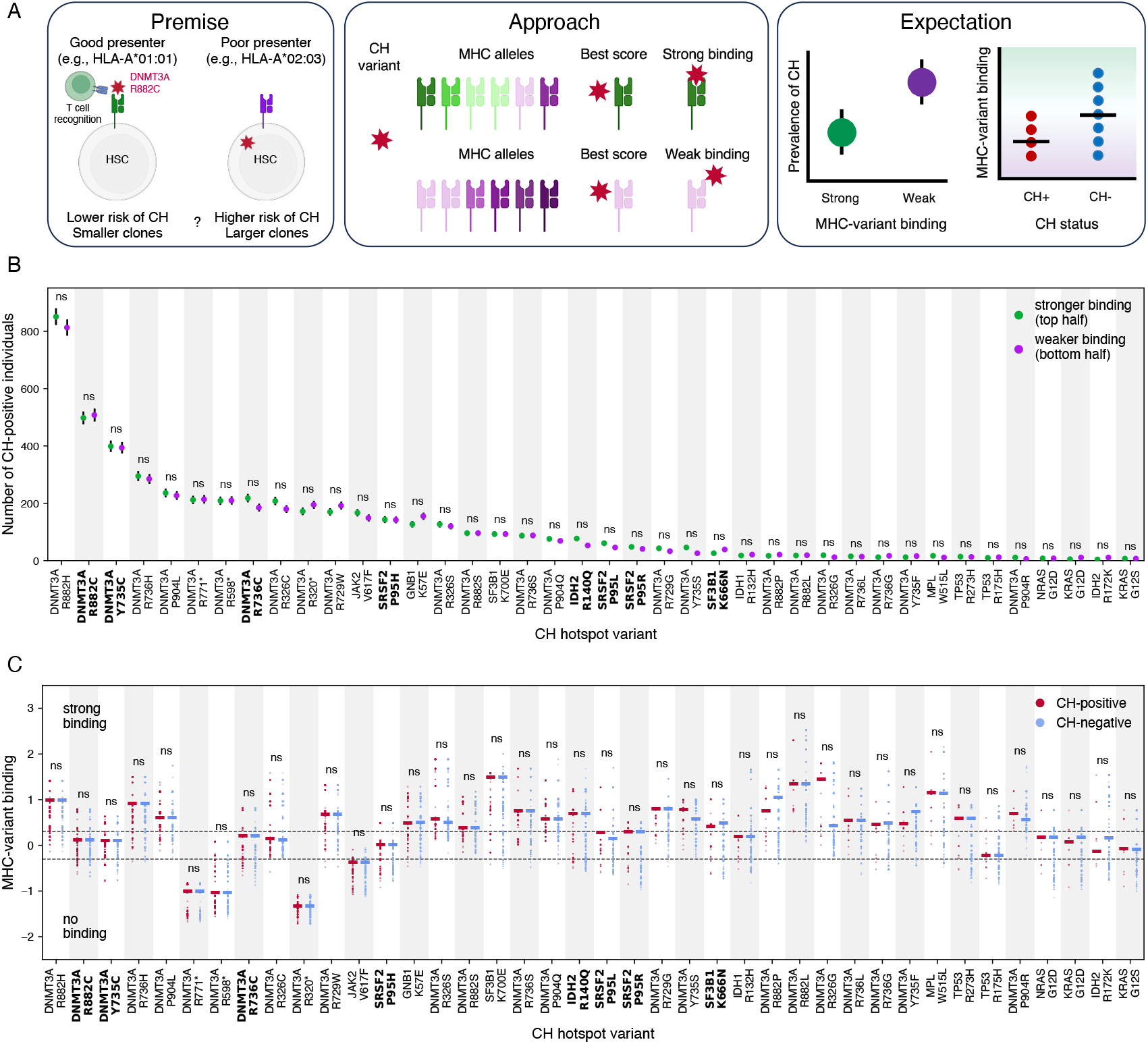
No effect of predicted MHC binding on CH prevalence. **A**. Overview of study hypothesis and approach. **B**. The number of CH-positive individuals does not differ between groups of varying relative MHC-variant binding. p-value from two-sided Fisher’s exact test (Bonferroni-corrected), ns - not significant. **C**. The distribution of MHC-variant binding scores does not differ between individuals who are positive and negative for each variant. For visualisation purposes, we compare the scores of all CH-positive individuals to 2,000 randomly sampled individuals who were identified as CH-negative for the variant. p-value from two-sided Mann Whitney U test (Bonferroni-corrected). Names of variants in bold indicate variants with at least 5 CH-positive individuals identified as binding the variant strongly and 5 who were predicted not to bind the variant. Dashed lines indicate threshold for strong binding (top line, corresponding to % elution rank 0.5) and weak binding (bottom line, % elution rank 2).

We identified 9, 309 individuals who carried a somatic variant in one or more of the hotspot positions using the UK Biobank exome sequencing dataset. First, we split the cohort into two equal-sized groups based on their relative predicted binding capacity and compared the number of CH carriers between the stronger- and weaker-binding groups (Fig. 1B). There was no significant difference in CH prevalence between stronger- and weaker-binding individuals for any of the variants studied. This analysis is powered to detect negative selective pressures between 0.5%-3% depending on the fitness of the variant and how many times it is detected (Supplementary Figure 3).

A different way of examining the same relationship is to ask whether the distribution of binding scores for a given variant differs between CH carriers and non-carriers. If MHC presentation prevents some variants from expanding, there would be an expectation that binding scores for individuals with expanded clones would fall towards the lower end of scores observed in the general population (Fig. 1A). We compared the distributions of scores between individuals positive and negative for each variant, noting that any clone at sufficiently high VAF to be observable in UK Biobank must have undergone a period of strong positive selection ^25^ (Fig. 1C). Again, we found no significant differences in predicted binding score between CH carriers and non-carriers across the 40 hotspots.

If binding to MHC requires affinities above a specific absolute threshold it is possible that examining absolute binding scores rather than relative binding would be a more effective way of detecting negative selection. We therefore sorted individuals using absolute (rather than relative) predicted binding score. For the 8 hotspots where at least five individuals carrying the variant were predicted to bind it strongly (% elution rank < 0.5) and at least five were predicted not to bind at all (% elution rank > 2), we compared CH prevalence between binding and non-binding individuals (Fig. 2). None of the 8 variants showed a significant difference in CH prevalence, suggesting again that MHC genotype is not influencing clonal expansions driven by these variants. Altering the binding threshold did not affect the results (Supplementary Fig. 5).

**Fig. 2.**
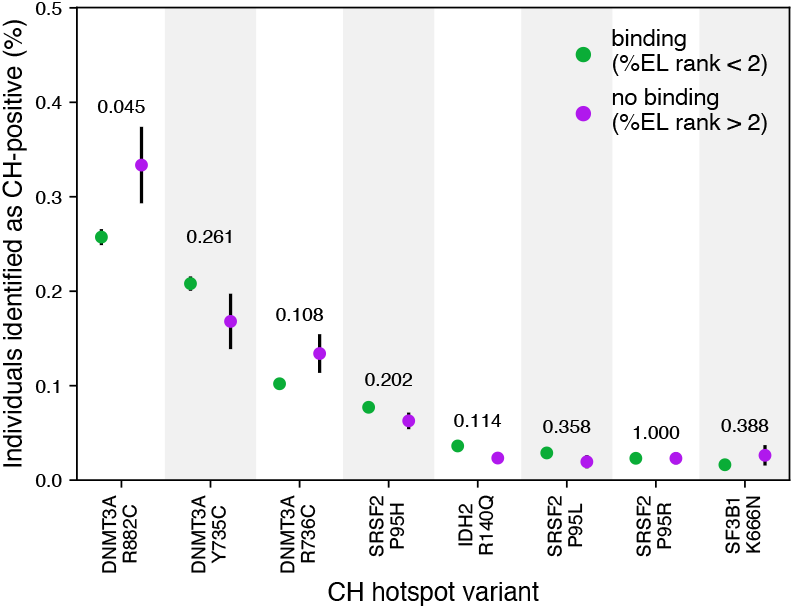
Effect of absolute binding on CH prevalence. Fraction of individuals identified as CH-positive between groups of different absolute bindings for the 8 variants where sufficient numbers of individuals carrying the variant were predicted to be strong binders (green data points) or non-binders (purple data points). p value from two-sided chi-squared test (Bonferroni corrected).

Recent studies have proposed an association between greater diversity in an individual’s MHC I and II alleles and protection against lung cancer ^26^ and some subtypes of lymphoma ^27^. Individuals with a more diverse MHC are able to present a greater diversity of mutant peptides, potentially conferring better protection against expansion of a range of mutant clones. Therefore, we tested if the number of MHC alleles and MHC heterozygosity status influence CH risk. However, we found no association between a greater diversity in MHC alleles and decreased prevalence of CH (Supplementary Fig. 6).

### Clone sizes unaffected by MHC binding

While the previous analysis suggests that comprehensive immune suppression of clonal expansion of HSCs driven by variants examined here is unlikely to occur in any significant fraction of the population, we may be able to detect more nuanced effects by examining the distribution of clone sizes. If immune surveillance was only initiated once clones reached a certain size, for example, then we would expect the clone size distributions for stronger- and weaker-binding individuals to diverge above a certain clone size (Fig. 3A). The clone size distribution in CH can be accurately estimated using the variant allele frequency (VAF) measured in peripheral blood. We divided carriers of each variant into equal-sized groups based on relative binding score and compared the distribution of VAFs between the stronger- and weaker-binding groups (Fig. 3B-J and Supplementary Fig. 7). We observed no significant difference between the stronger- and weaker-binding VAF distributions either on an individual variant level (Fig. 3C-J and Supplementary Fig. 7) or with all variants aggregated together to increase power (Fig. 3B). In this analysis, we are powered to detect negative selective pressures of 4% for variants detected in ∼100 individuals, 2.5% for variants detected in ∼300 individuals, and 1% for variants detected in 1000 individuals (Supplementary Fig. 8).

**Fig. 3.**
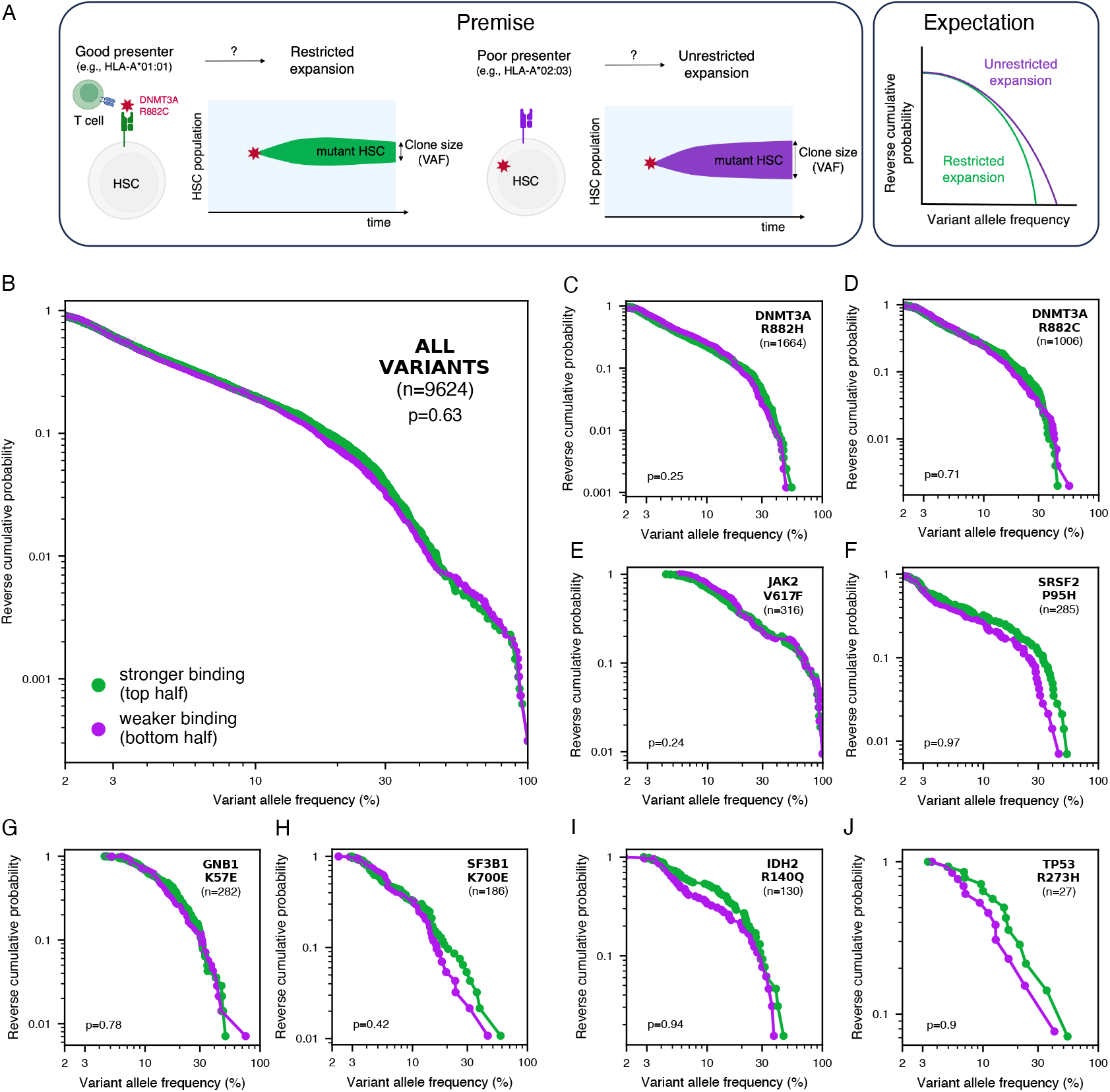
No effect of predicted MHC binding on clone size distribution. **A**. Premise and expectation. **B**. The distribution of clone sizes is not associated with differences in clone sizes across all variants. **C–J**. Relationship between predicted binding capacity for several representative CH variants. p-value from Kolmogorov-Smirnov test (one-sided).

As above, we repeated this analysis splitting the cohort according to absolute predicted binding score, comparing individuals predicted to bind the mutant peptide against those predicted not to bind at all. We found no significant differences in clone size distributions between these groups (Fig. 4 and Supplementary Fig. 9).

**Fig. 4.**
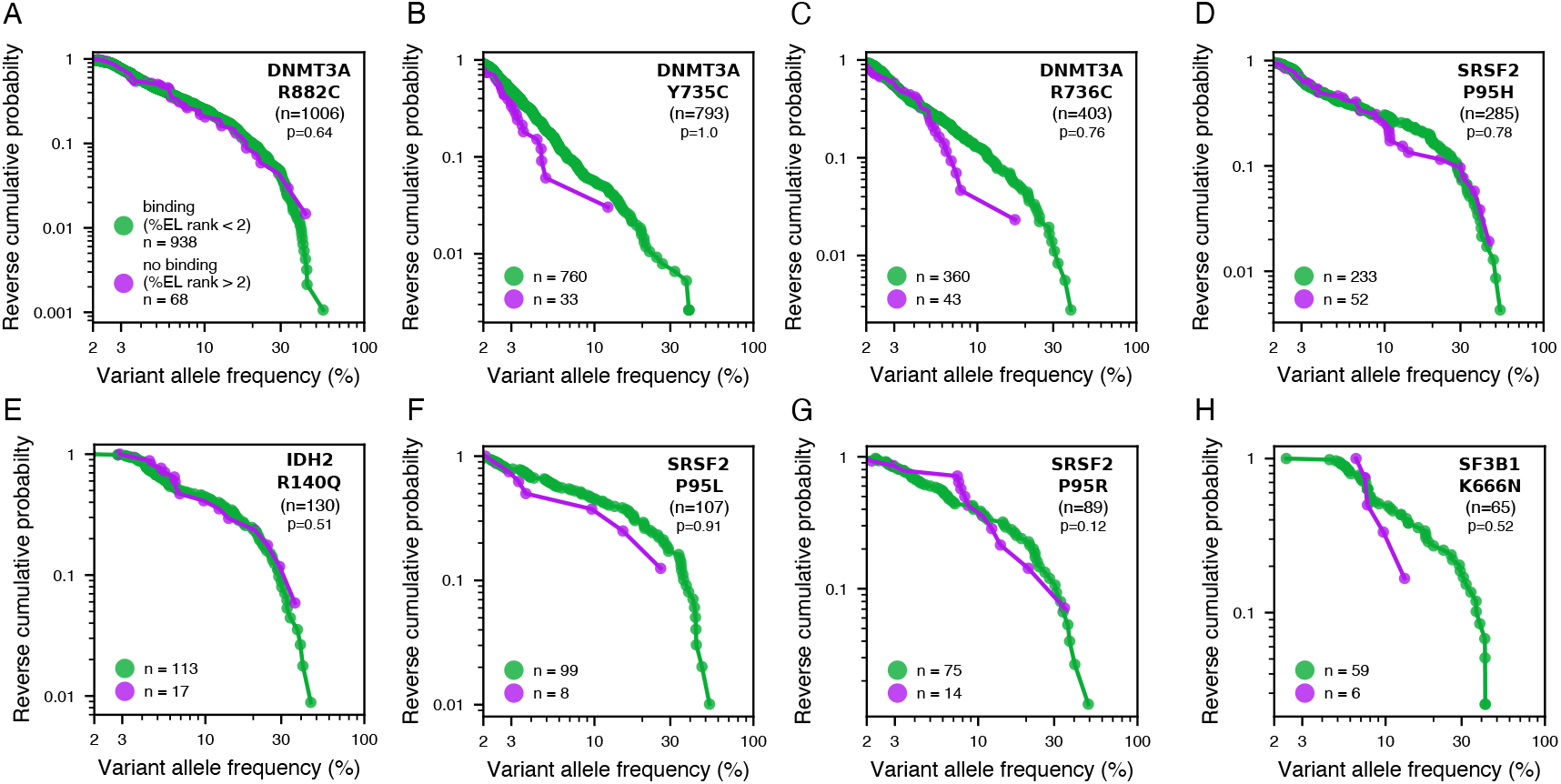
No effect of predicted MHC binding on clone size distribution. **A-H**. Reverse cumulative clone size distributions for strong predicted MHC binders (green) and predicted non-binders (purple) based on absolute binding thresholds. p-value from Kolmogorov-Smirnov test (one-sided).

HSCs have been shown to express MHC II class alleles ^8,28^, and this was suggested to allow surveillance of HSCs by CD4+ T cells and elimination of cells carrying mutations that generate immunogenic peptides, providing protection from cancer ^6,8^. Therefore, we also tested the association with binding capacity defined by MHC II genotype, using predictions obtained from *NetMHCIIpan-4*.*3* ^29^. However, we saw no evidence for an association between MHC II genotype and risk and progression of CH driven by specific variants (Supplementary Fig. 14-17).

Our analysis relies on accurate predictions of MHC-peptide binding, therefore, to mitigate against possible biases associated with the specific prediction software, we also used predictions of variant binding to MHC I from a different method, PRIME2.0^30^. However, in all cases we reached the same conclusions (Supplementary Fig. 10-13).

### Variants with functionally validated MHC-restrictions

A limitation of our previous analysis is the reliance on computation prediction of MHC binding. Given that MHC binding predictions do not directly correspond to immunogenicity, we asked if we could observe signs of immune surveillance of variants which are known to be MHC-restricted (i.e., presented only on specific MHC alleles). Were a given variant to be well presented on a specific MHC allele and capable of eliciting an immune response, the expectation would be that this allele would be under-represented in individuals affected by CH driven by the variant compared to the general population. Therefore, we selected variants where there is experimental support for their MHC restriction (KRAS G12D ^31,32^, IDH1 R132H ^33^, IDH2 R140Q ^34,35^, TP53 R175H ^36,37^), and compared the frequency of MHC alleles known to present those variants between the UKB cohort and individuals positive for those variants. We also tested JAK2 V617F in HLA-B*35:01 and HLA-A*02:01 backgrounds, as these has been previously reported to be under-represented in MPN patients carrying the JAK2 mutation ^38^. However, we did not find evidence for those alleles to be less common in individuals in whom the variant expanded than in the UKB cohort (Fig. 5), as would have been expected if MHC presentation led to strong immune surveillance.

**Fig. 5.**
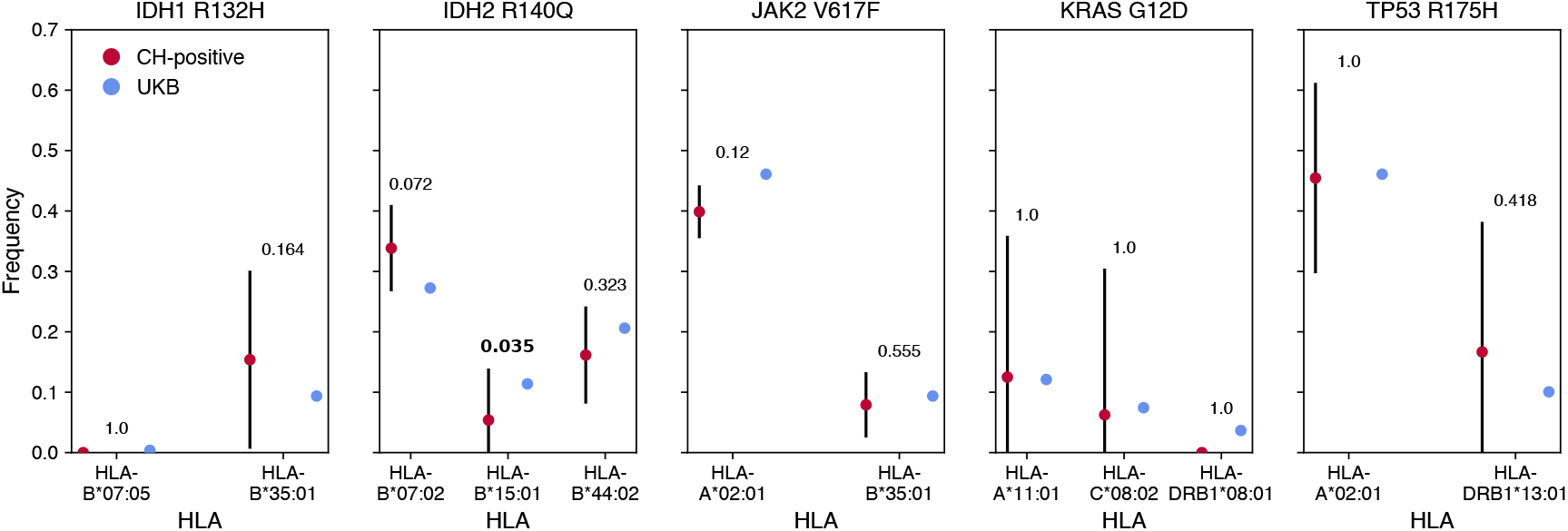
Limited evidence for effect of MHC genotype in MHC-restricted immunogenic variants. Comparison of the frequencies of the MHC alleles experimentally validated to present specific CH hotspot variants between individuals harbouring the variant and the general UKB population. p-value from Fisher’s exact test.

## DISCUSSION

Here, we have attempted to shed light on the question of whether the immune system exerts negative selective pressures on clones that arise and expand during somatic evolution in ageing blood by examining the association between the MHC genotype and variants which successfully expand in CH. Despite the scale of the UKB affording considerable power to detect weak selective pressures, we found no evidence that the MHC-based presentation plays a role in selecting which clones successfully expand in a given individual, arguing against a significant impact of adaptive immune surveillance on somatic evolution of the blood. We did not detect a difference in predicted MHC binding capacity between individuals who were CH positive or negative for any of the variants examined, and did not observe differences in proportions of CH-affected individuals across groups of different predicted binding capacity to any variant. Moreover, we did not observe a consistent relationship between predicted binding capacity and the distribution of clone sizes in individuals positive for a given variant. Therefore, whilst absence of evidence is not evidence of absence, our results would argue against strong effect of immune surveillance in shaping the somatic diversity of blood.

Our results are consistent with studies which did not identify immune response in somatic tissues despite a significant burden of mutations in somatic clones ^17,18^ and analyses suggesting that selective pressure exerted by the immune system is stronger in cancer than pre-cancerous lesions ^39,40^. A possible reason for this would be could be the requirement for clones carrying non-self antigens to reach a threshold fraction for T-cell recognition ^20,41^. For the range of clone sizes examined here, we do not find evidence for a threshold clone size being required for immune recognition in CH, though deeper error-corrected sequencing might be required to detect such effects.

Our analysis relies on computational predictions of MHC binding, the accuracy of which may be imperfect ^42^ and which do not directly correspond to peptide immunogenicity ^43–46^, therefore it is possible that some of the variants examined here would not be sufficiently immunogenic to induce an effective immune response even if predicted to be strongly bound by the MHC. This could explain why some other mutations, such as CALR frameshifts which can drive myeloproliferative neoplasms, have been argued to be under early immune surveillance based on the presence of memory T cells and antibodies to CALR in healthy individuals ^47^. It is possible that immune surveillance could occur early, but only for highly immunogenic mutations, which also seem to trigger immune response at lower threshold of clonal fraction ^20^. We note that in our study, we have not seen evidence of surveillance for variants where there is experimental evidence that they are immunogenic on certain MHC backgrounds, e.g., KRAS G12D ^48^, TP53 R175H ^36^, IDH2 R140Q ^34^ and IDH1 R132H ^49^. However, we cannot exclude that clones which expand successfully evade immune detection through other mechanisms which cannot be established based on the available data alone, such as reduction in MHC expression or immunosuppressive environment in the bone marrow. For example, JAK2 V617F-positive clones were suggested to downregulate MHC ^38^ and increase expression of PD-L1^50^, which could explain why despite immunogenicity, only minor responses to JAK2 V617F were found in MPN patients ^51^. As MHC II expression levels are lower in hematopoietic progenitor cells than HSCs, which cell type the clonal expansion originated from could reduce the effectiveness of immune surveillance of mutant progenitor cells ^8^. Research suggests these mutations which drive clonal expansions in CH in the elderly can be acquired as early as in utero ^52–55^. As this developmental stage is associated with a more tolerogenic immune environment ^56^, mutations acquired early in life could possibly be tolerated, such that expansions of these clones later in life would not be under surveillance. It is also possible that immune recognition has occurred, but due to the slow nature of the clonal expansions in CH, that T-cells become exhausted due to persistent exposure to the antigen ^57^.

Previous studies suggested that individuals’ predicted MHC binding to mutations present in tumours is poorer than for mutations absent from the tumours, suggesting that tumour development occurs in the ‘gaps’ in an individual’s immune system ^5,9^. We reproduced this analysis for CH by comparing each individual’s predicted binding score for the CH variant they carry (‘present’) with their respective scores for the remaining common CH mutations *not* observed in that individual (‘absent’) (Supplementary Fig. 18). While the average binding scores for CH carriers varied considerably between variants, we did not observe consistently lower binding scores between the present CH driver and the other possible (absent) mutations. Aggregating these scores across variants can lead to apparent significant differences in the binding capacity of present versus absent variants (Supplementary Fig. 19). However, this apparent effect is driven predominantly by the MHC binding of the most common variants. In this case this effect is due to the variants which are most common (DNMT3A R882 mutations) also being predicted to be bound universally well, and would be observed even if all individuals presented the variant equally well ^58^. Our observation that the variation in binding a given variant between individuals is lower than variation in binding capacity between variants would suggest that in the case of many driver mutations, MHC presentation may not have major effects on which of them are allowed to expand, and therefore the effect may not be as robust as previously suggested ^5,9^. We note that the UKB cohort is rather uniform with respect to its ancestry and genetic make-up, and we may see larger variation in MHC binding in more diverse cohorts ^59^, and that there may be cases of mutations which generate peptides which are both strongly immunogenic and sufficiently differentially bound between MHC alleles where such an effect would still be observed ^6^, but it may not generalize across all cases of driver mutations.

CH is known to be associated with increased risk of hematological cancers ^21,60^ and cardiovascular disease ^61^, therefore limiting expansions of CH clones, e.g., through vaccination, would be an attractive strategy to reduce the risk of associated diseases. While our analysis suggests that immune surveillance does not affect the expansions of most known drivers of CH, it does not exclude that an immune response could be elicited in this way. Functional studies would be needed to evaluate the possibility of altering the evolutionary trajectory of clones expanded in CH through such interventions.

## METHODS

### UK Biobank exome sequencing

Exome sequencing of 450,000 people was performed as part of the UK Biobank, ^62,63^, a prospective cohort study of 500,000 middleaged people in the UK. All UKB participants signed a written informed consent form at enrolment. Ethical approval was given by the North West Multi-Centre Research Ethics Committee (REC 21/NW/0157).

### MHC typing in the UK Biobank

MHC types were imputed by Bycroft et al. at two-field (four-digit) resolution for 11 classical HLA genes (HLA-A, HLA-B, HLA-C, HLA-DRB1, HLA-DRB3, HLA-DRB4, HLA-DRB5, HLA-DQA1, HLA-DQB1, HLA-DPA1 and HLA-DPB1) using the HLA*IMP:02 algorithm with a multi-population reference panel ^23^. The MHC genotyping dataset contains posterior probability values for each MHC allele (0 – not present, 1 – one allele present, 2 – both alleles present). According to published recommendation (^64^, a threshold of 0.7 was used to identify individuals positive for a given allele, and 1.5 to identify individuals carrying two copies of a given allele. Individuals were excluded from further analysis if not all of their alleles could be genotyped with confidence, filtering was done separately for MHC I and MHC II genotype to avoid excluding individuals who only had MHC I or MHC II genotype missing.

### Calling CH hotspot variants from UK Biobank exome sequencing

CH hotspots were called from the UK Biobank exome CRAM files using the DNAnexus Research Analysis Platform (application 28126) ^65^. *samtools mpileup* (docker: *quay*.*io/biocontainers/samtools:1*.*12–hd5e65b6_0*) was used to identify variants at 26 genomic positions in 11 genes, corresponding to common single nucleotide variants identified as potential drivers of CH by Watson et al. ^25^ (Supplementary table 1). As part of a separate analysis, variants in TP53 were obtained from the same CRAM files using *mutect2*, a somatic variant caller available in the Genome Analysis Toolkit (docker: *broadinstitute/gatk:4*.*1*.*3*.*0*) ^66^. We restricted our analysis to variants which were present in more than 15 individuals in the entire UKB cohort. The number of unique variants analysed (after all filters were applied) was 40.

To remove false positives caused by sequencing errors, variants not supported by at least 2 variant reads were discarded (3 in TP53). Individuals carrying a mosaic chromosomal alteration overlapping the variant (as identified by ^67^) were excluded. Since certain kinds of systemic cancer treatment are known to alter the selection landscape of CH ^68^, individuals with a history of cancer before blood draw were excluded, including self-reported cancers (with the exception of basal cell carcinoma, pre-cancer of the cervix and rodent ulcer).

After the CH and MHC filtering steps, there were 384, 600 individuals with MHC I allele data available of whom 9, 309 carried one or more CH variants (9, 624 variants). For MHC II there were 327, 961 individuals and 7, 932 individuals (8, 212 variants).

### MHC-variant binding predictions

The genomic coordinates of CH variants were obtained from COSMIC ^69^. Corresponding amino acid sequences were obtained from the UniProt database (The UniProt Consortium, 2023). We obtained peptide sequences containing the position where the mutation occurs of appropriate length (8-11 for MHC class I and 15 for MHC class II).

Predictions were generated for all MHC alleles typed at least once in the UKB which were available in NetMHC-pan4.1^24^ or NetMHCIIpan-4.3^29^. This included 194 MHC class I alleles (A type, B type, C type) and 441 MHC class II alleles or allele combinations (DRB1, combinations of DPA-DPB alleles and combinations of DQA-DQB alleles). DRB3, DRB4 and DRB5 alleles were excluded from analysis as they were missing in 50% of the examined population.

To obtain predictions for each variant-allele combination, we first generated binding predictions for each variant-carrying peptide and allele combination and chose score of the best-scoring peptide as the variant score. % minimum elution rank score (%EL rank) was used in the main analysis as it is considered the most biologically relevant metric.

For each individual, we obtained a single score for each CH variant by identifying the best (lowest) %EL rank across combinations of the gene with all alleles carried by this individual. We obtained two separate scores, one for MHC class I and the other for MHC class II.

### Logistic regression analysis

For all examined individuals in whom MHC genotype were typed with high confidence, we defined MHC heterozygosity status for each locus (MHC class I: A, B, C; MHC class II: DPA, DPB, DQA, DQB, DRB1) and number of distinct MHC I and MHC II alleles typed. We also collected data on sex, age at first assessment and smoking status (1 - ever smoker, 0 - not smoking). Due to missing data, we excluded individuals of non-European ancestry. For each individual, we defined CH diagnosis as presence of 2 or more reads corresponding to at least one CH variants examined here (3 in case of variants in TP53). Individuals were excluded from regression analysis if age data was missing.

### Power analysis

To assess the power of our study to detect differences in variant prevalence and VAF distributions between cohorts, we performed simulations using a custom Python script adapted from Watson et al. ^25^. We simulated clonal dynamics for a range of fitness coefficients (0.1 to 0.2) and sampled different numbers of individuals to approximate the distributions observed in the UK Biobank. At each level of sampling we determined the minimum difference in fitness effect for which the prevalence was distinguishable using the Fisher’s exact test at a significance level of 0.05, and for which the VAF distributions were distinguishable using the Kolmogorov-Smirnov test at a significance level of 0.05. We sampled each number of individuals five times to assess the extent of variability in the power to identify differences between different fitness effects.

## Data availability

The analysis of UK Biobank data was conducted using the UK Biobank Resource under Application Number 28126. None of the funders had a role in data analysis, decision to publish or preparation of the manuscript.

## Code availability

Code used for the analysis is available on GitHub: https://github.com/the-blundell-lab/MHC_ClonalHaematopoiesis

## Acknowledgments

We thank all members of the Blundell lab for input. This research has been conducted using the UKB Resource under application number 28126. J.R.B. is funded by a UKRI Future Leaders Fellowship and by the CRUK Cambridge Cancer Centre. J.R.B, B.W. and H.A.J.M are supported by the International Alliance for Cancer Early Detection, an alliance between Cancer Research UK (C14478/A29329), Canary Center at Stanford University, the University of Cambridge, OHSU Knight Cancer Institute, University College London and the University of Manchester. Schematic diagrams for figures 1A and 3A were created using BioRender.com

## Author contributions

J.R.B conceived of the study. J.R.B. and B.W. designed the study with input from H.A.J.M. All analysis was developed by B.W and J.R.B. with input from H.A.J.M. Manuscript was written by B.W. and edited by all authors.

