## supplementary figures for "No evidence of immunosurveillance in mutation-hotspot driven clonal haematopoiesis"

#### Supplementary Table

**Supplementary table 1.** Putative CH drivers targeted in the analysis, originally identified by Watson et al.[25]. Genomic positions refer to assembly build 38 (GRCh38).

| Gene | Target variant(s) | chr:position |
|---|---|---|
| DNMT3A | R320* | 2:25247647 |
| | R326C/G/S | 2:25247629 |
| | R598* | 2:25244214 |
| | R729G/W | 2:25240439 |
| | Y735C/F | 2:25240420 |
| | R736H/L | 2:25240417 |
| | R736C/G/S | 2:25240418 |
| | R771* | 2:25240313 |
| | R882C/S | 2:25234374 |
| | R882H/L/P | 2:25234373 |
| | W860R | 2:25235727 |
| | P904L/Q/R | 2:25234307 |
| GNB1 | K57E | 1:1815790 |
| IDH1 | R132H | 2:208248388 |
| IDH2 | R140Q | 15:90088702 |
| | R172K | 90088606 |
| JAK2 | V617F | 9:5073770 |
| KIT | D816H/Y | 4:54733154 |
| | D816V/F | 4:54733155 |
| KRAS | G12D/V | 12:25245350 |
| | G12C | 12:25245351 |
| MPL | W515L | 1:43349338 |
| NRAS | G12D/V | 1:114716126 |
| | G12C | 1:114716127 |
| SF3B1 | K666N | 2:197402635 |
| | K700E | 2:197402110 |
| SRSF2 | P95H/R/L | 17:76736877 |

#### Supplementary Figures

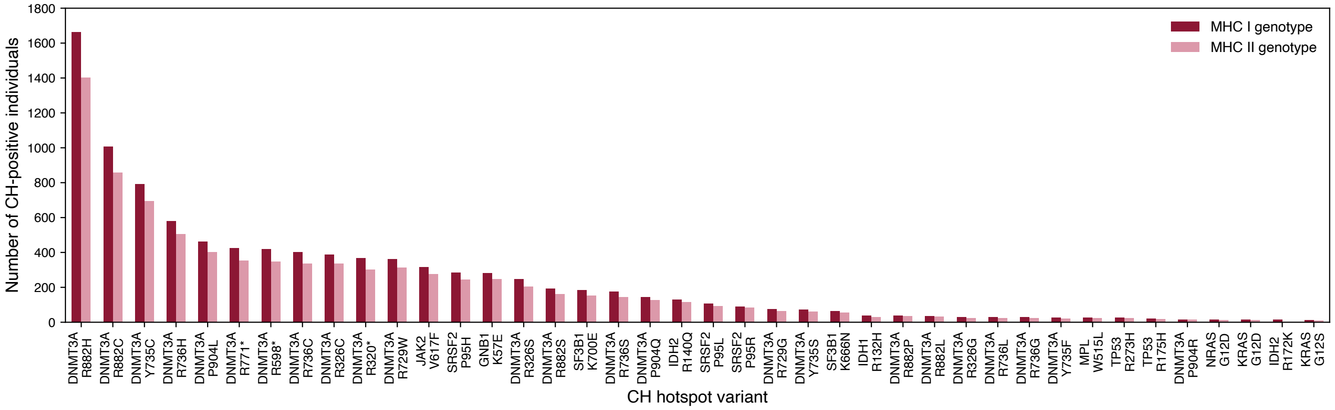


**Extended data Fig. 1. CH hotspot variants in the UKB** Number of individuals carrying each variant identified in the UKB with complete MHC I or MHC II genotype data.

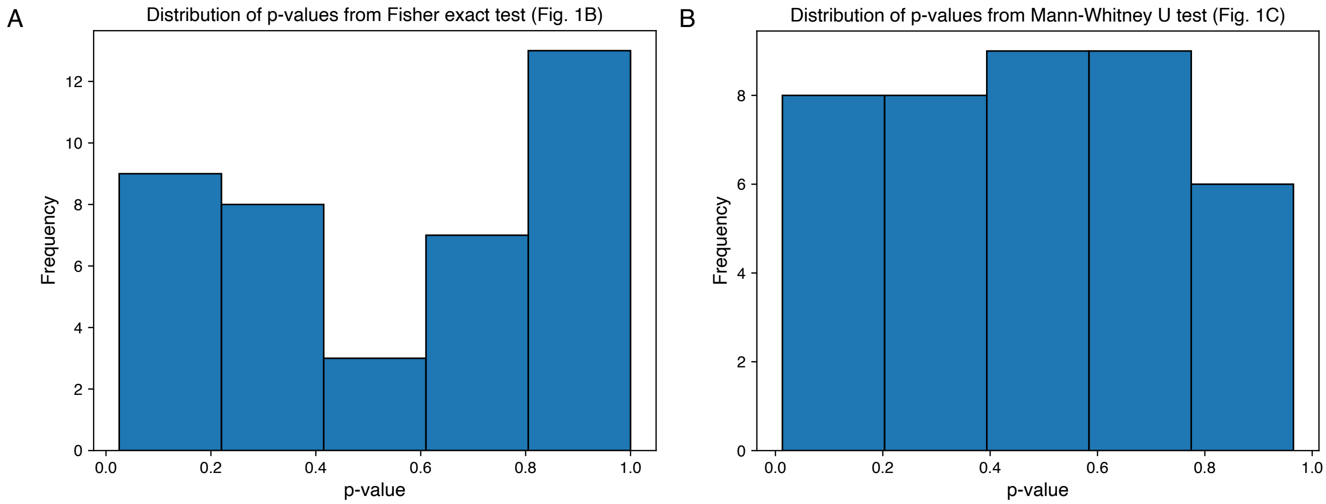


**Extended data Fig. 2. Statistical analysis A.** Distribution of p-values from Fisher's exact test conducted on Fig. 1B to analyse the differences in the number of CH-positive individuals between better and worse binding groups for each CH variant. **B.** Distribution of p-values from Mann Whitney U test conducted on Fig. 1C to compare the MHC-variant binding scores between CH-positive and CH-negative individuals.

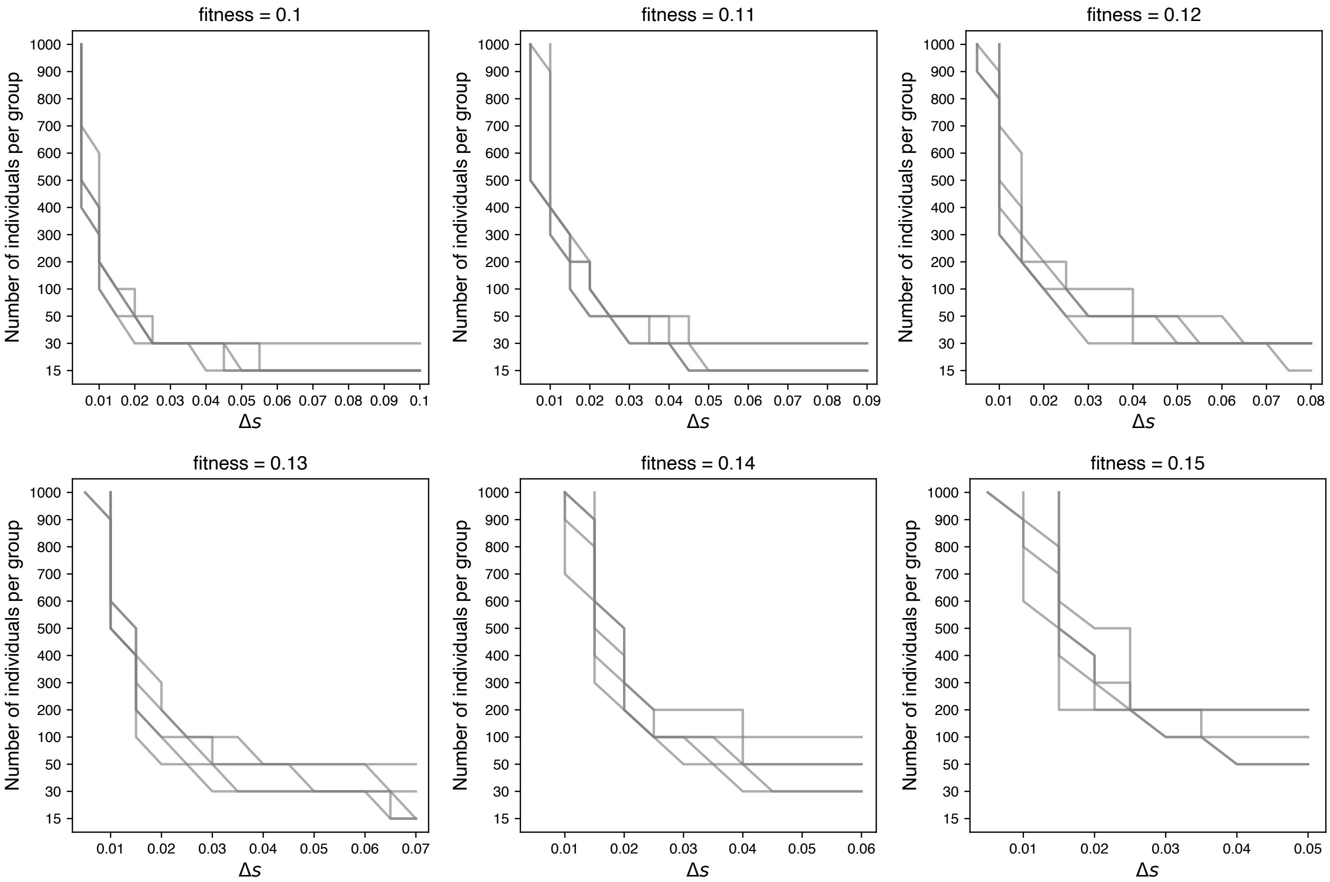


**Extended data Fig. 3. Power analysis** Results of simulations to determine extent of power to detect the difference in the Fisher's exact test. Grey lines indicate the boundary between significant and non-significant results for independent sampling of number of individuals. Related to Fig. 1B.

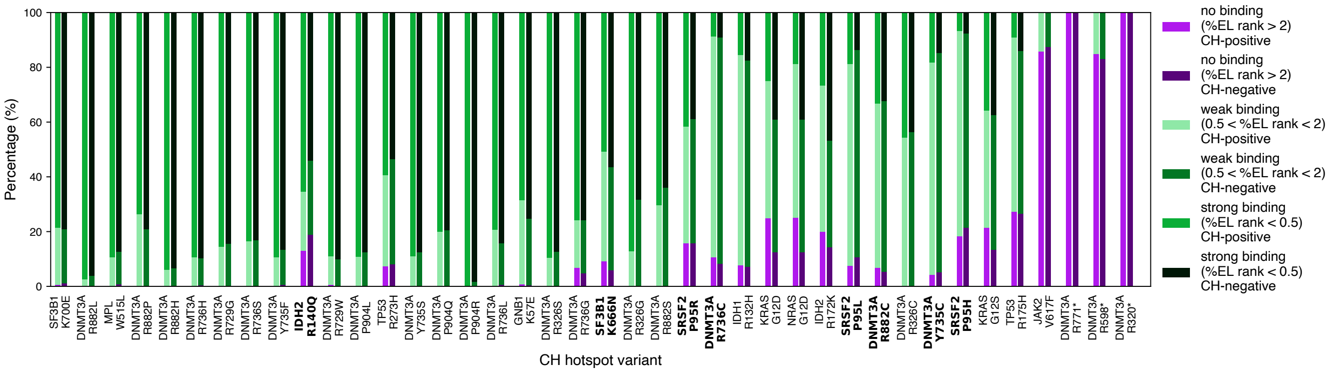


**Extended data Fig. 4. No difference in percentage of the population predicted to bind a given variant between CH-positive and CH-negative individuals** Distribution of individuals who are predicted to bind each variant strongly, weakly or not based on their MHC I genotype, in CH-positive and CH-negative individuals for each variant. Names of variants in bold indicate variants with at least 5 CH-positive individuals identified as binding the variant strongly and 5 who were predicted not to bind the variant.

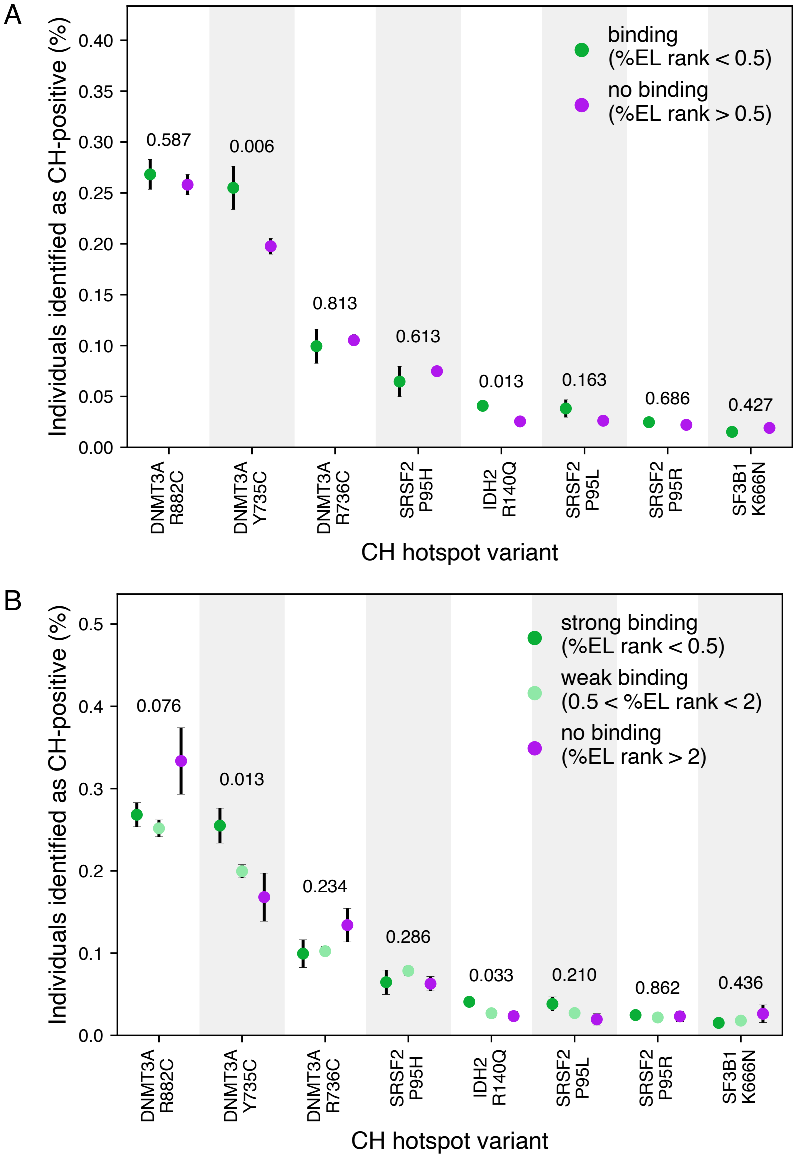


**Extended data Fig. 5.** **Effect of MHC-variant absolute binding on CH prevalence**. **A.** Comparison of the fraction of CH-positive individuals present in groups predicted to bind vs not bind the variant. **B.** Comparison of the fraction of CH-positive individuals present in groups predicted to bind the variant strongly, weakly vs not bind it. p-value from chi-squared test. Related to Fig. 2.

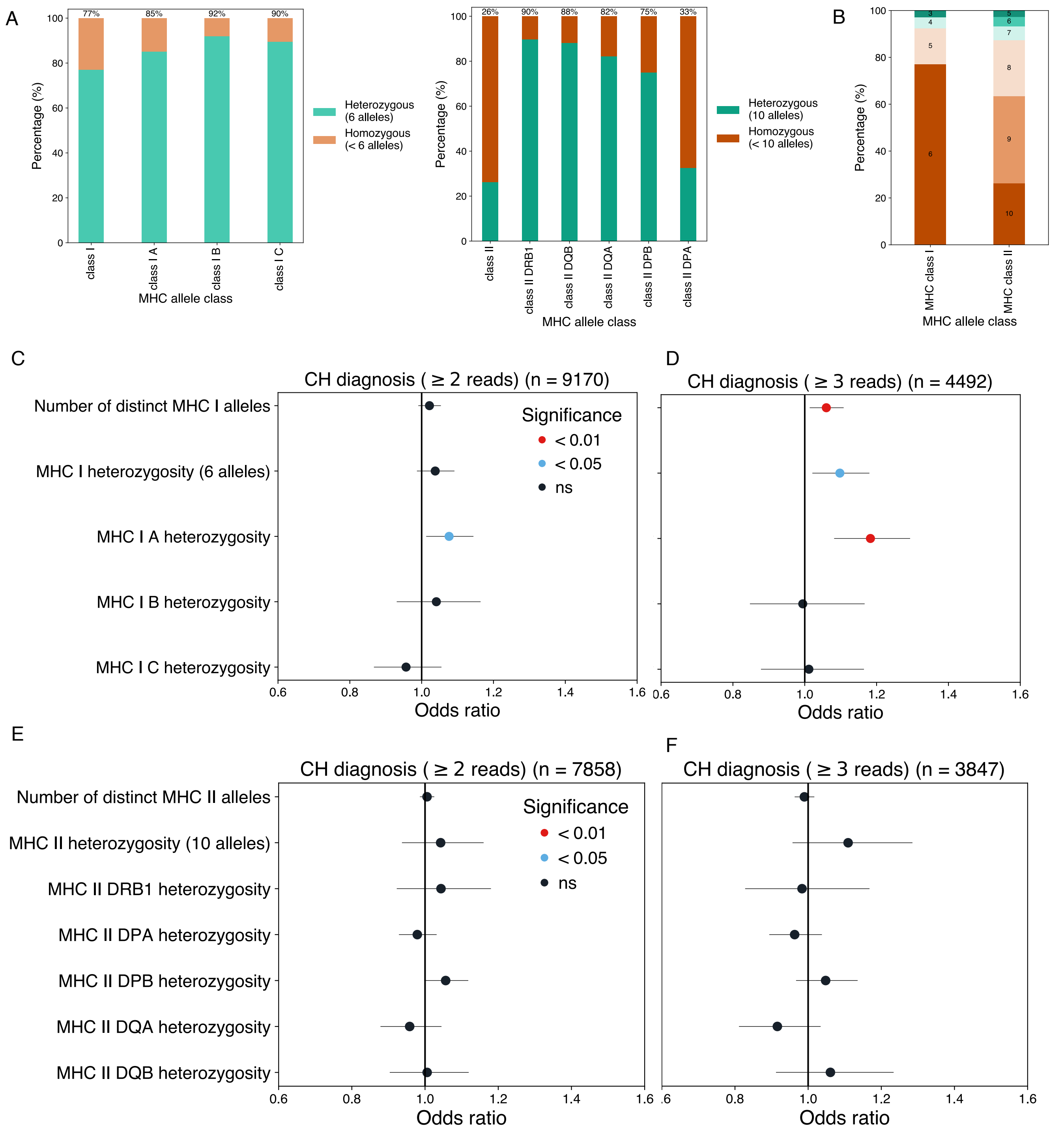


**Extended data Fig. 6. No impact of heterozygosity status on overall risk of CH. A.** Distribution of individuals who are heterozygous vs homozygous for each major class of MHC I or MHC II alleles in the UKB cohort. **B.** Distribution of the number of MHC I and MHC II alleles in the UKB cohort. **C-D** Relationship between number of MHC I alleles, MHC I heterozygosity status (overall vs specific alleles) and CH diagnosis with threshold of 2 (C.) or 3 (D.) reads. **E.-F.** Relationship between number of MHC I alleles, MHC I heterozygosity status (overall vs specific alleles) and CH diagnosis with threshold of 2 (E.) or 3 (F.) reads.

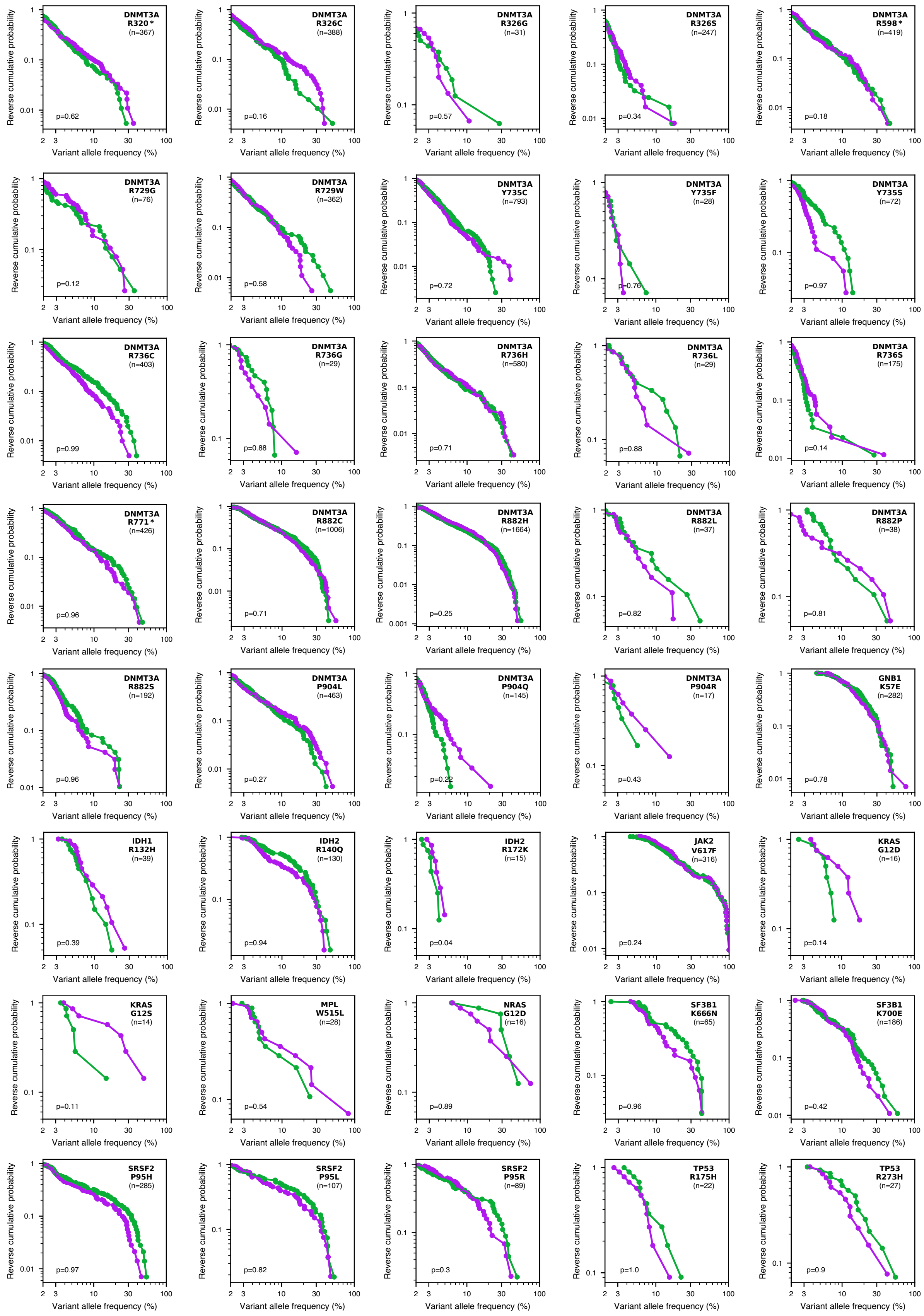


**Extended data Fig. 7. Relationship between predicted MHC-variant binding capacity and clone size in CH-positive individuals for each of the variants examined.** p-value from Kolmogorov-Smirnov test (one-sided). Related to Fig. 3.

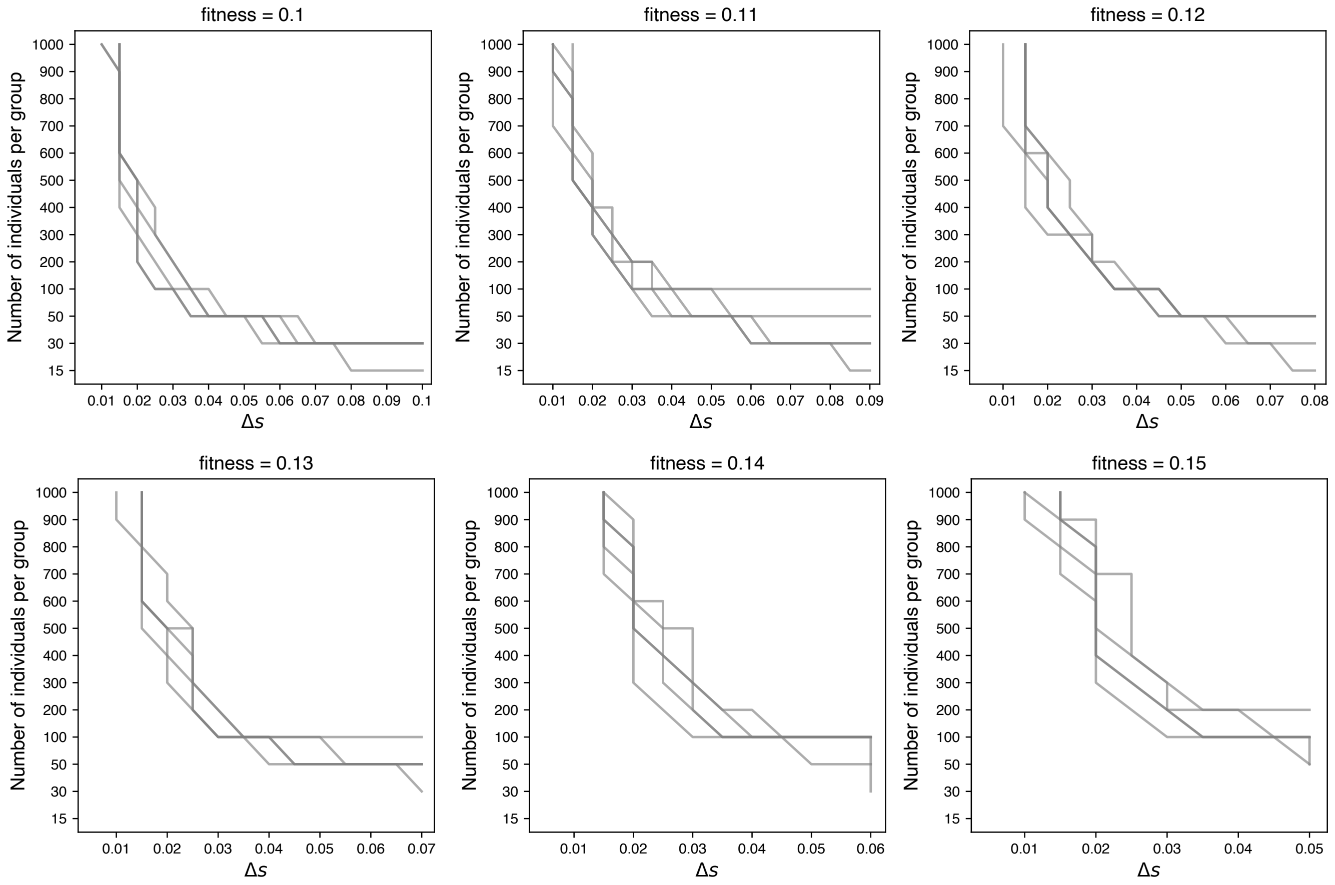


**Extended data Fig. 8. Power analysis.** Results of simulation to determine extent of power to detect the difference in clone sizes in Kolmogorov-Smirnov test. Grey lines indicate the boundary between significant and non-significant results for independent simulations. Related to Fig. 3 and Supplementary Fig. 7.

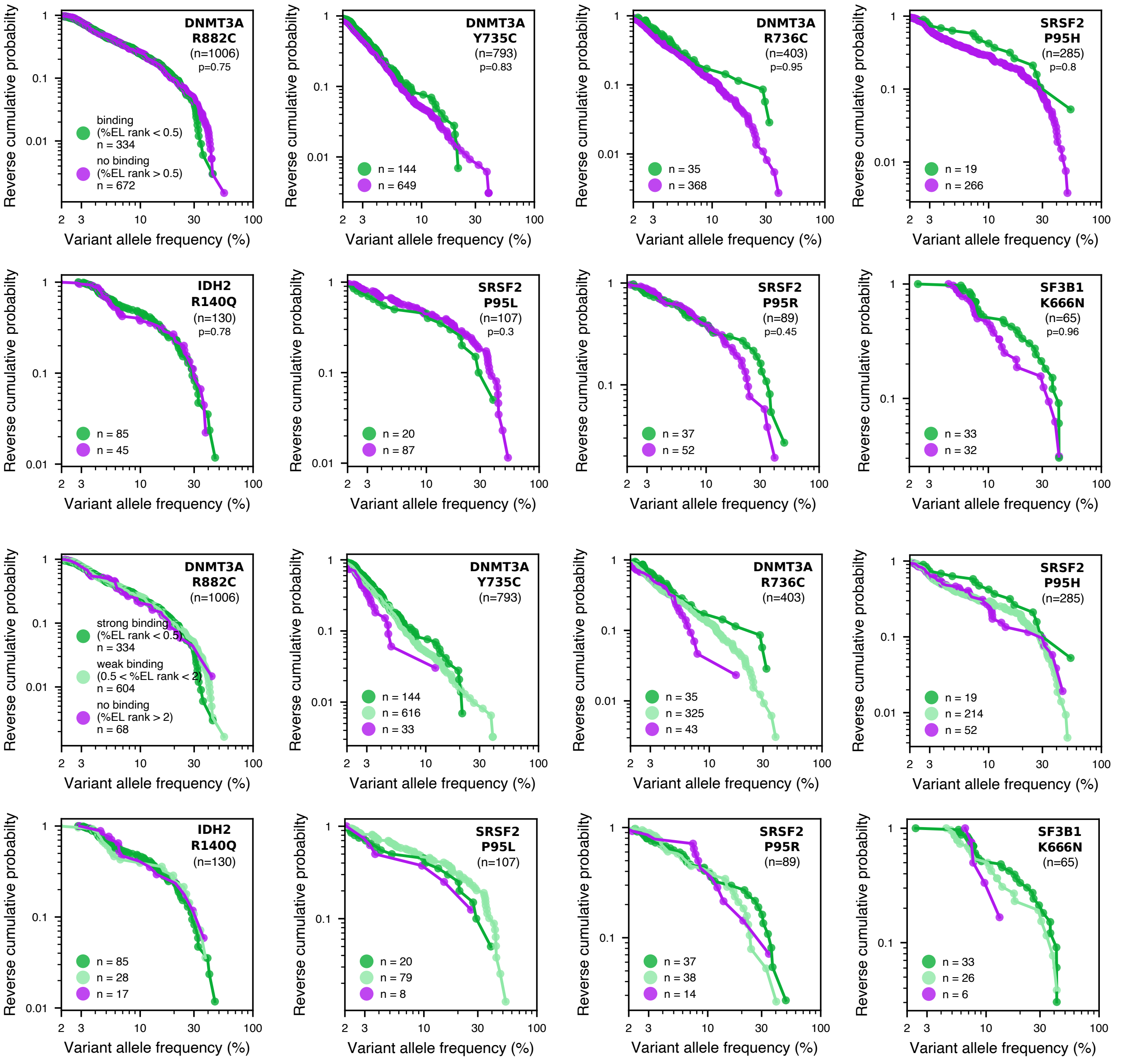


**Extended data Fig. 9.** **Effect of predicted MHC binding on clone size distribution.** Absolute binding assessed at threshold of % elution rank 0.5 (strong binding) and at two thresholds (% elution rank 0.5 and % elution rank 2). p-value from Kolmogorov-Smirnov test (one-sided). Related to Fig. 4.

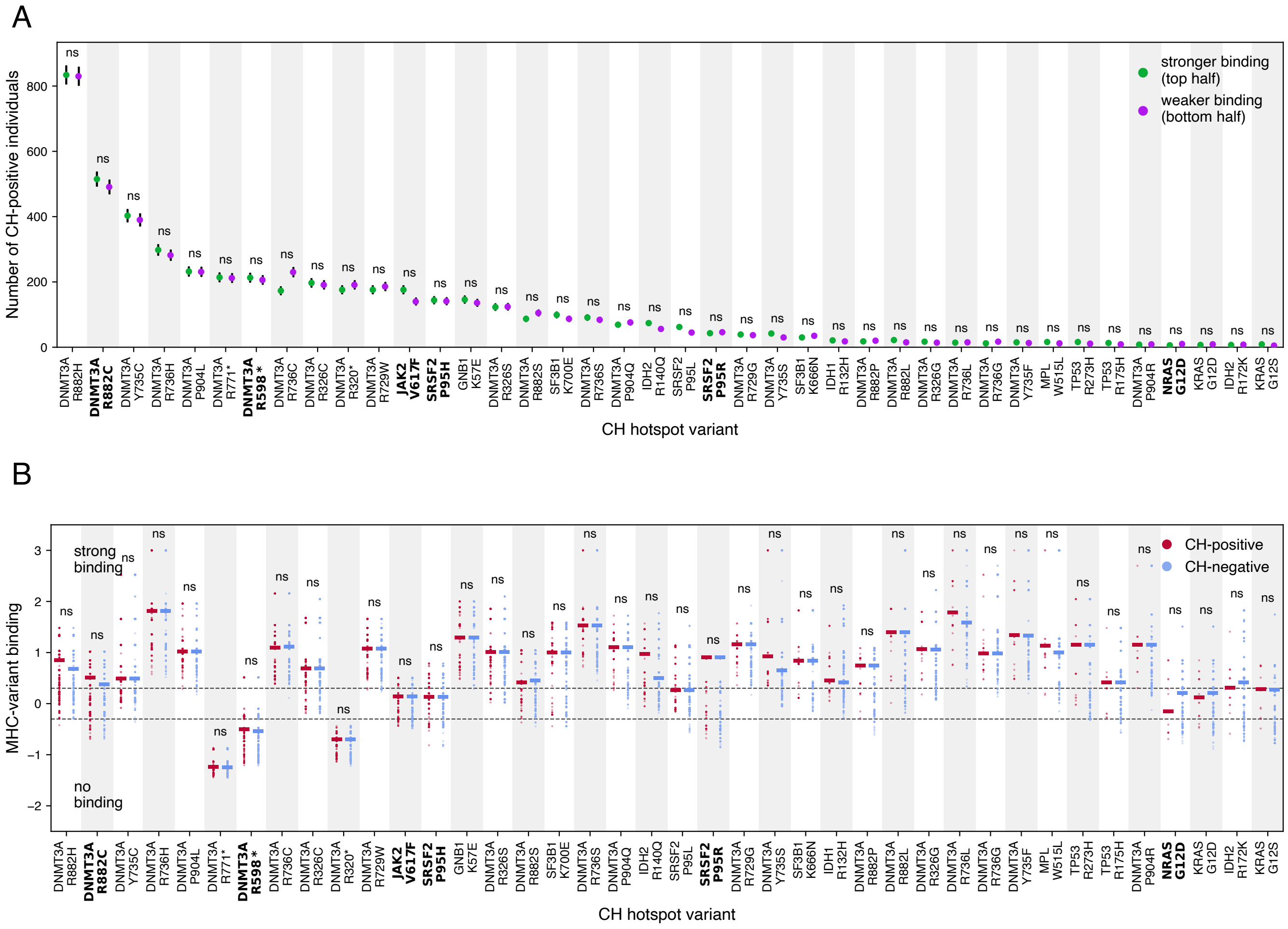


**Extended data Fig. 10. Additional analysis based on binding predictions from PRIME2.0. A.** Comparison of the number of CH-positive individuals between equal-sized groups predicted to bind a given variant better or worse. **B.** The distribution of MHC-variant binding scores between CH-positive and 2,000 randomly sampled CH-negative individuals. Variants with at least 5 CH-positive individuals identified as binding the variant strongly and 5 who were predicted not to bind the variant. Dashed lines indicate threshold for strong binding (top line) and weak binding (bottom line). Star indicates significance at 0.05 (Bonferroni-corrected), ns - not significant. Names of variants in bold indicate variants with at least 5 CH-positive individuals identified as strongly binding the variant and 5 who were predicted not to bind the variant. Related to Fig. 1.

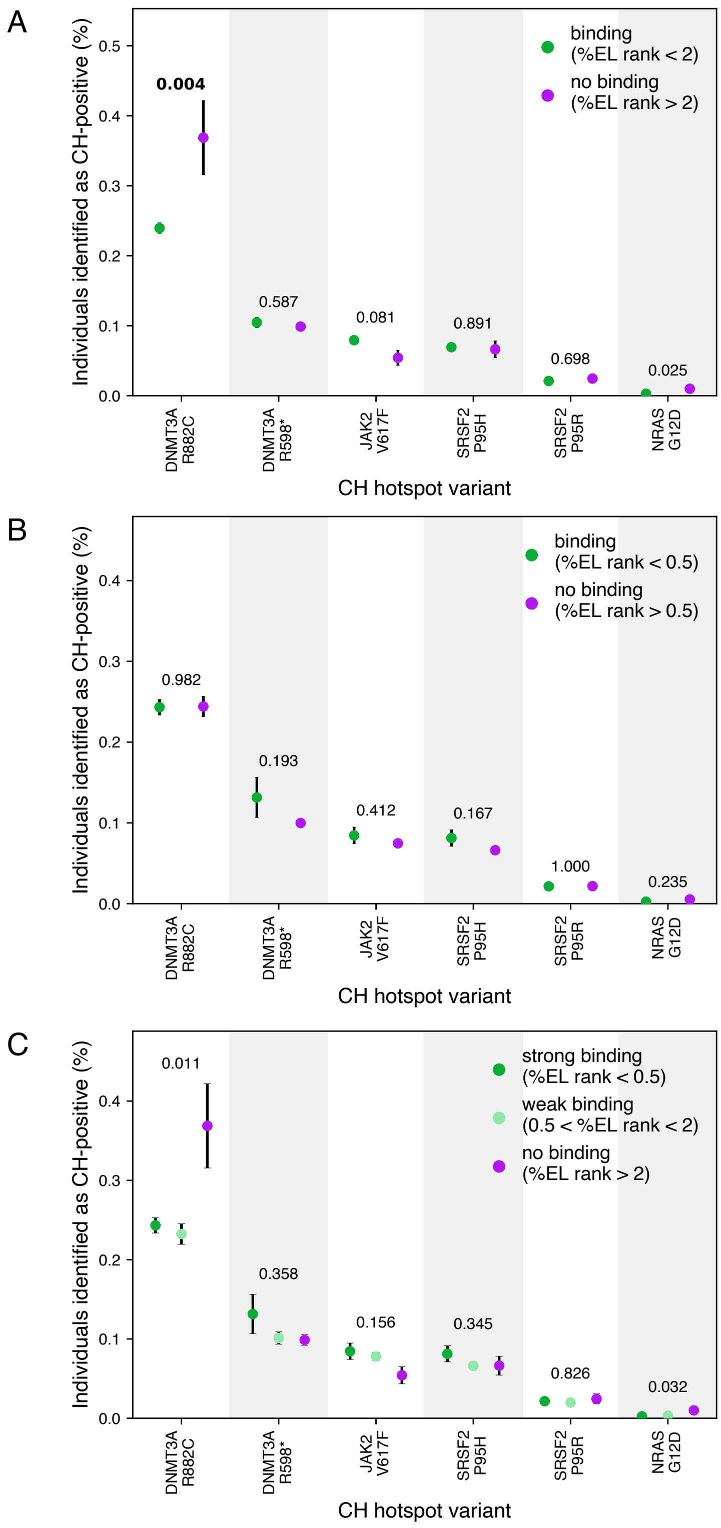


**Extended data Fig. 11. Additional analysis based on binding predictions from PRIME2.0**[30] **A.** Comparison of the fraction of CH-positive individuals present in groups predicted to be able to bind the variant (strongly or weakly) vs not bind it. **B.** Comparison of the fraction of CH-positive individuals present in groups predicted to bind strongly vs bind it weakly or not. **C** Comparison of the fraction of CH-positive individuals present in groups predicted to bind the variant strongly, weakly vs not bind it. p-value from chi-squared test. Related to Fig. 2 and Supplementary Fig. 5.

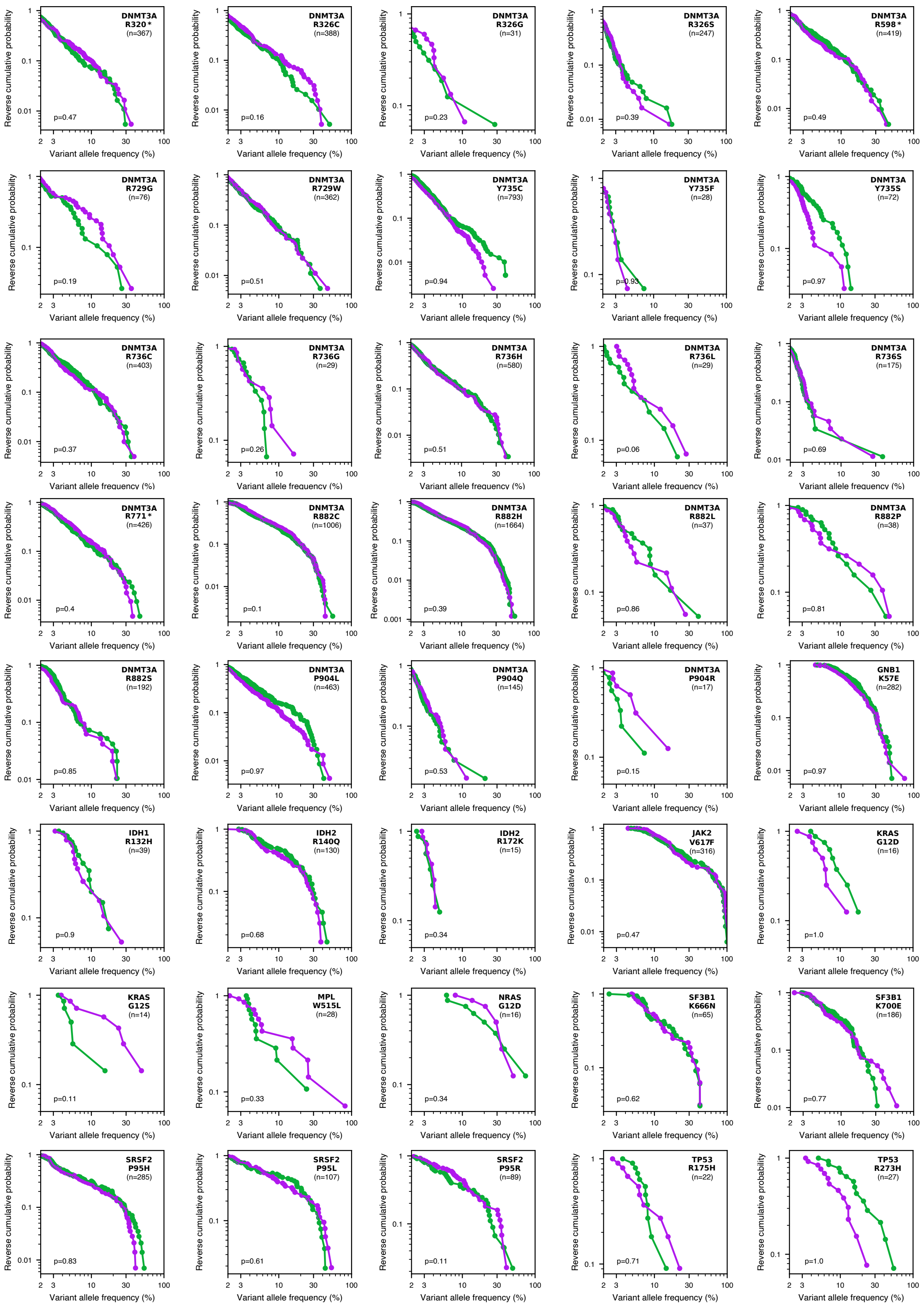


**Extended data Fig. 12. Additional analysis based on binding predictions from PRIME2.0.** Relationship between predicted MHC-variant binding capacity and clone size in CH-positive individuals for each of the 40 variants examined. p-value from Kolmogorov-Smirnov test (one-sided). Related to Fig. 3 and Supplementary Fig. 6.

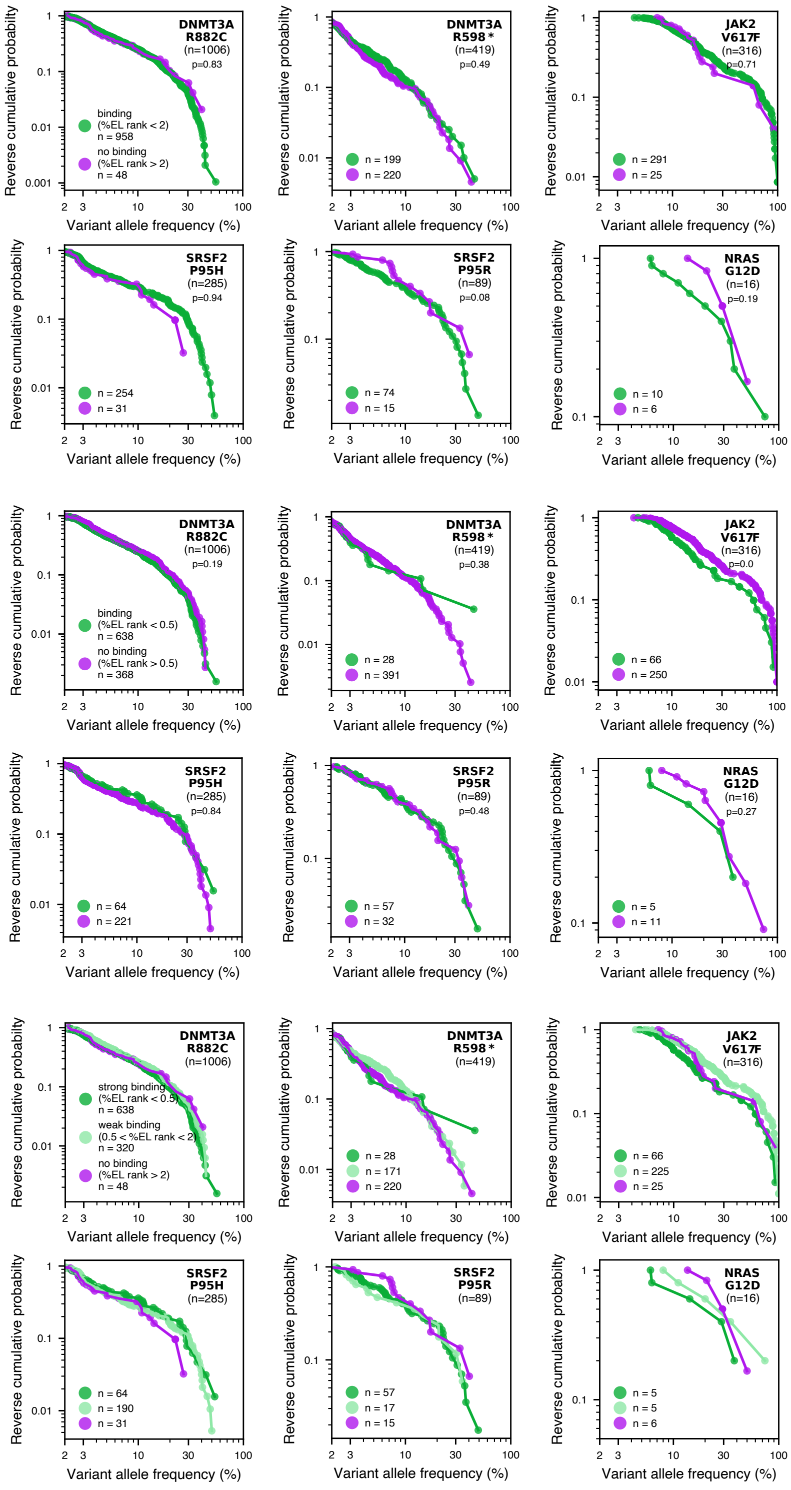


**Extended data Fig. 13. Additional analysis based on binding predictions from PRIME2.0.** Comparison of reverse cumulative distributions between CH-positive individuals who were classified as binding vs non-binding. p-value from Kolmogorov-Smirnov test (one-sided). Related to Fig. 4.

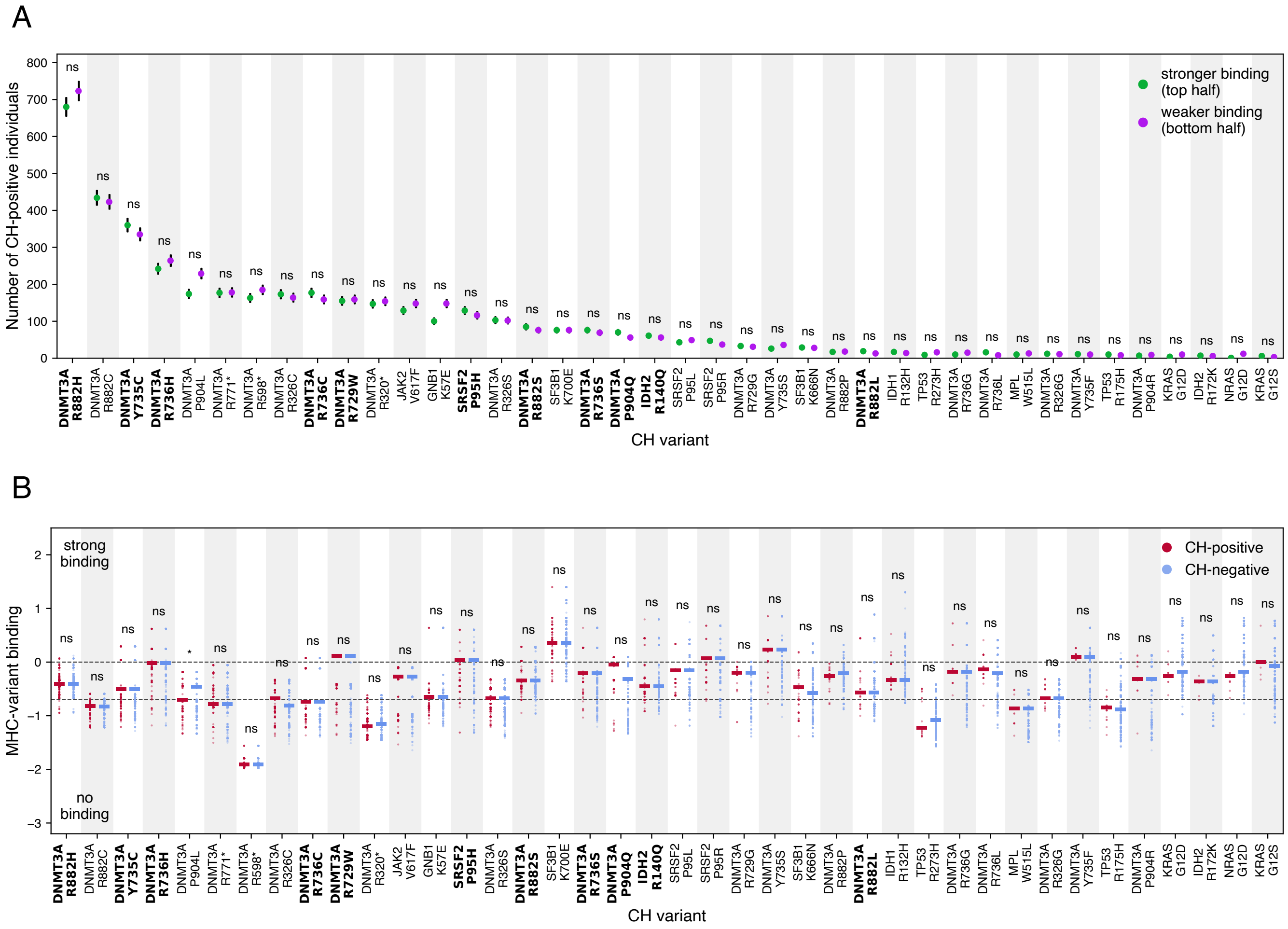


**Extended data Fig. 14. Additional analysis based on binding predictions between CH variants and MHC II alleles**. **A.** Comparison of the number of CH-positive individuals between equal-sized groups predicted to bind a given variant better or worse based on their MHC II genotype. p-value from Fisher's exact test (Bonferroni-corrected). **B.** The distribution of MHC-variant binding scores between CH-positive and 2,000 randomly sampled CH-negative individuals. Variants with at least 5 CH-positive individuals identified as strongly binding the variant and 5 who were predicted not to bind the variant. Dashed lines indicate threshold for strong binding (top line) and weak binding (bottom line). p-value from Mann-Whitney U test (Bonferroni-corrected). Names of variants in bold indicate variants with at least 5 CH-positive individuals identified as strongly binding the variant and 5 who were predicted not to bind the variant. Related to Fig. 1.

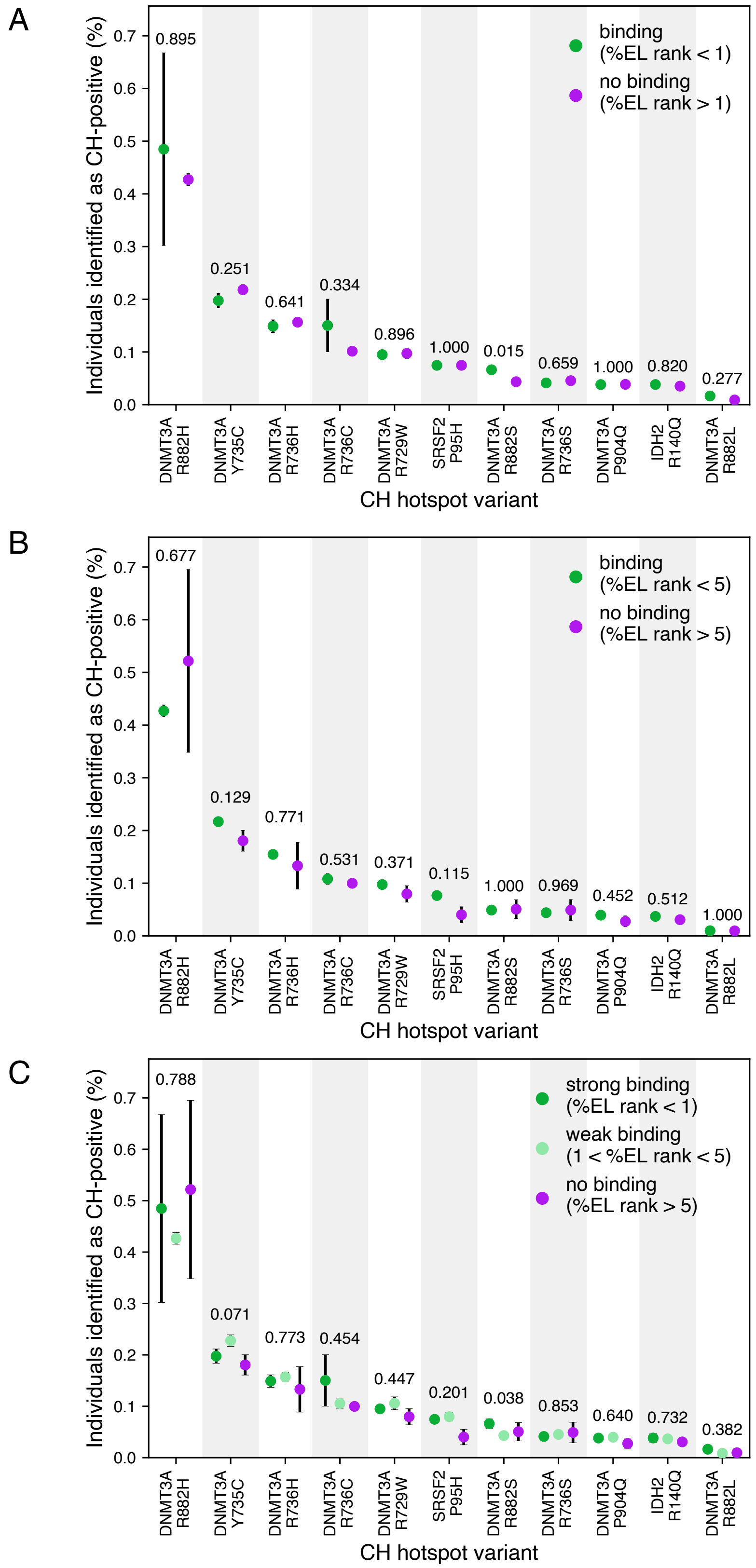


**Extended data Fig. 15. Additional analysis based on binding predictions between CH variants and MHC II alleles**. Fraction of individuals identified as CH-positive between groups predicted to have differential capacity of variant binding based on MHC II genotype. **A.** Comparison of the fraction of CH-positive individuals present in groups predicted to bind vs not bind the variant (threshold %EL rank = 5). **B.** Comparison of the fraction of CH-positive individuals present in groups predicted to bind vs not bind the variant (threshold %EL rank = 1). **C.** Comparison of the fraction of CH-positive individuals present in groups predicted to bind the variant strongly (threshold %EL rank = 1.), weakly vs not bind it. p-value from Chi-squared test. In bold, indicated p-value significant after multiple testing correction. Related to Fig. 2

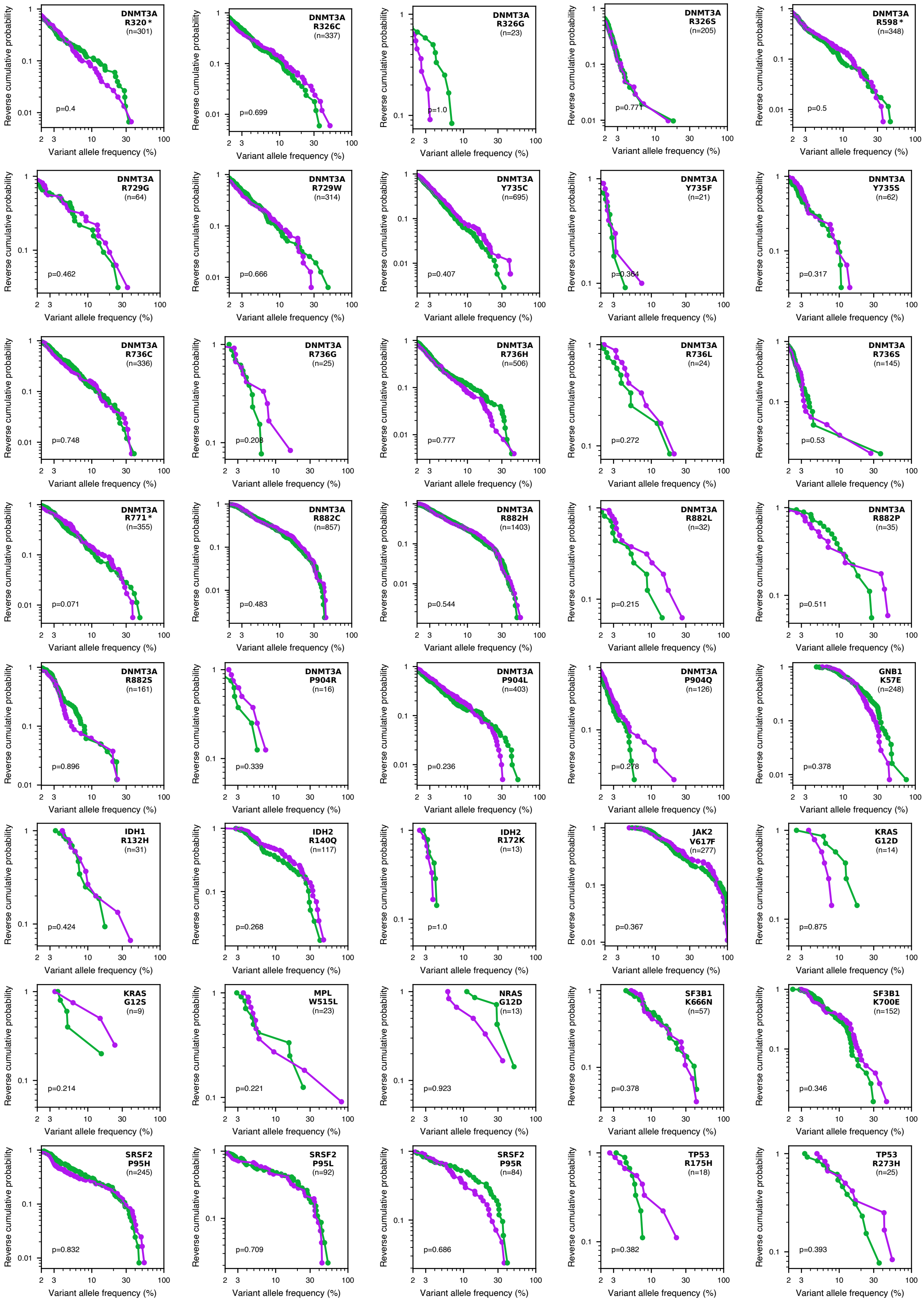


**Extended data Fig. 16. Additional analysis based on binding predictions between CH variants and MHC II alleles.** Comparison of reverse cumulative distributions between CH-positive individuals predicted to be in top vs bottom halves of binding each examined variant based on MHC II genotype. p-values from Kolmogorov-Smirnov test (one-sided). Related to Fig. 3.

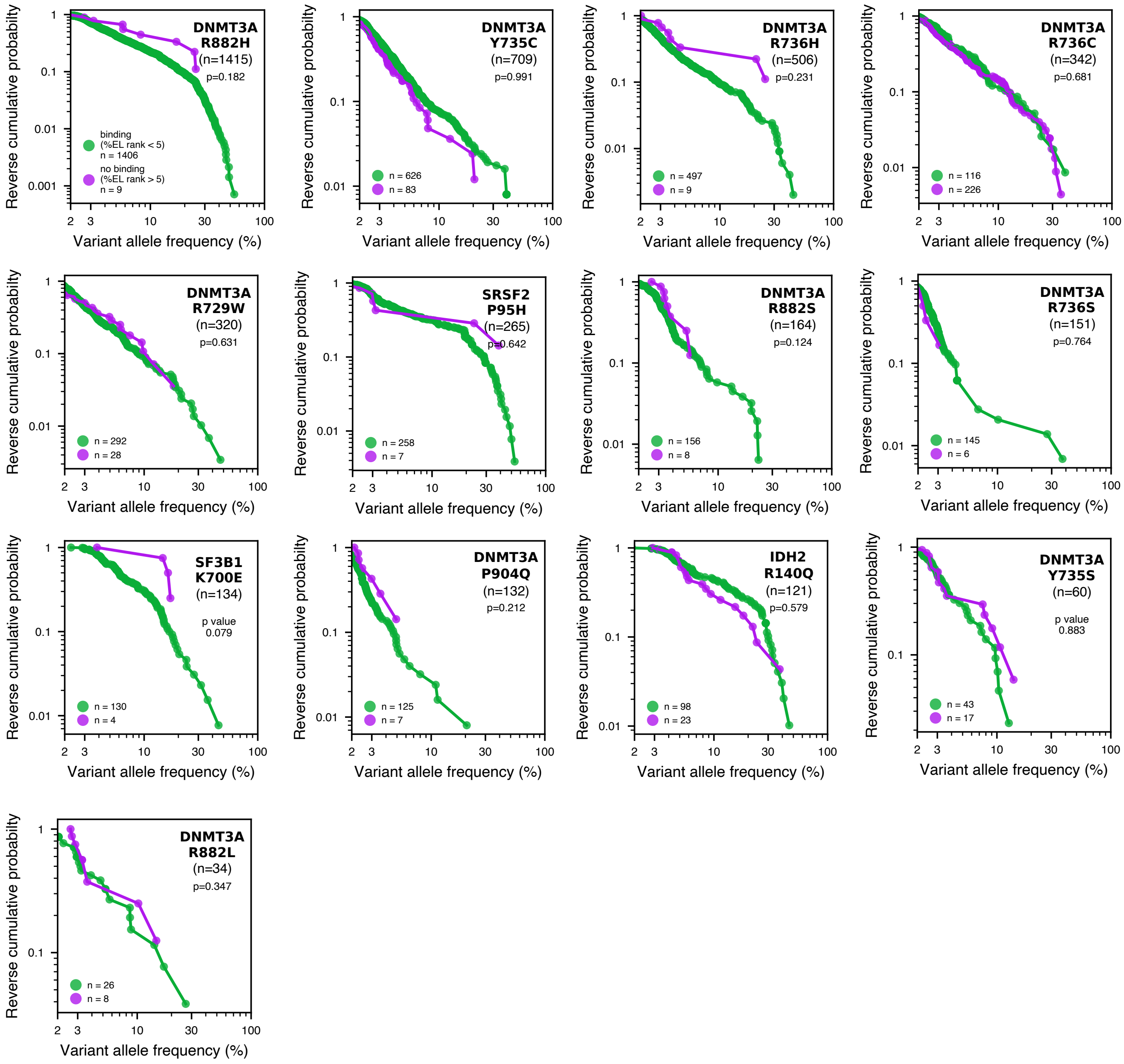


**Extended data Fig. 17. Additional analysis based on binding predictions between CH variants and MHC II alleles**. Comparison of reverse cumulative distributions between CH-positive individuals who were classified as binding vs non-binding based on MHC II genotype. p-values from Kolmogorov-Smirnov test (one-sided). Related to Fig 4.

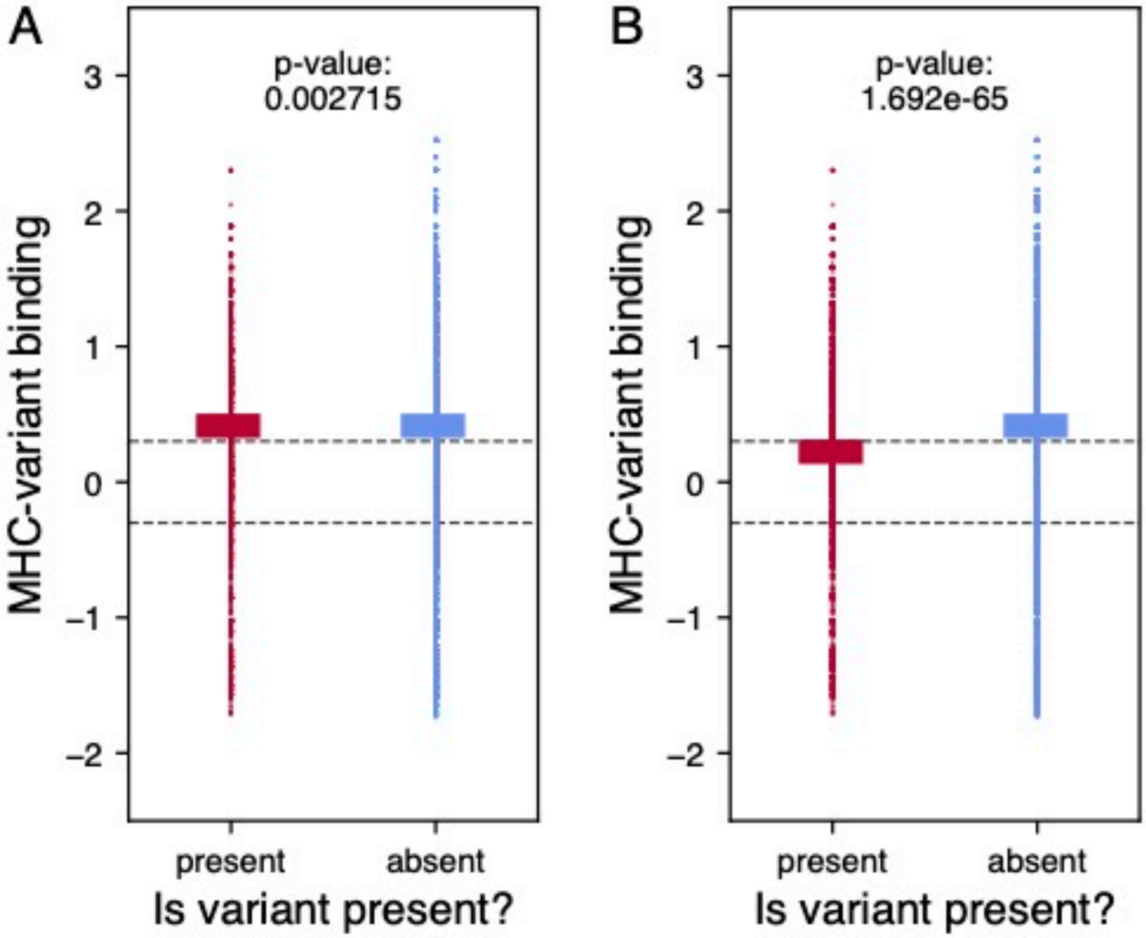


**Extended data Fig. 18. Binding predictions vary significantly more between variants than between individuals for a given variant** Distribution of scores for variant identified in the individual ('present') and the other 39 possible driver variants examined ('absent'). Analysis carried out on individuals who carry only one, rather than multiple, variants.

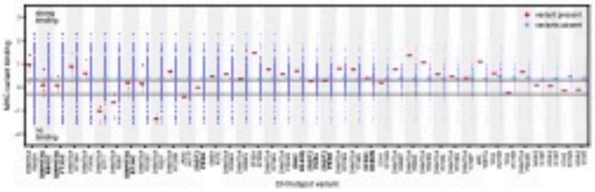

**Extended data Fig. 19. Comparison of scores for present vs absent variants when aggregated**. **A.** All variants (40) included. **B.**. Two most common variants, DNMT3A R882C and DNMT3A R882H, removed from analysis. p-values from Mann-Whitney U test. Related to Supplementary Fig. 18.

### SUPPLEMENTARY MATERIAL

#### A. Supplementary Figures.


- Supplementary figure 1 Number of CH-positive individual with confidently genotyped MHC I and MHC II for each variant
- Supplementary figure 2 Distribution of p-values from Fisher's exact test and Mann-Whitney U test presented on Fig. 1.
- Supplementary figure 3 Analysis of power in the Fisher's exact test (Figure 1)
- Supplementary figure 4 Distribution of individuals with different binding scores across 40 variants tested
- Supplementary figure 5 Absolute MHC-variant binding vs CH prevalence at different %elution rank thresholds
- Supplementary figure 6 Analysis of MHC heterozygosity vs CH risk
- Supplementary figure 7 Relative MHC-variant binding vs CH clone size across 40 variants tested
- Supplementary figure 8 Analysis of power in the Kolmogorov-Smirnov test (Figure 3)
- Supplementary figure 9 Absolute MHC-variant binding vs CH clone size at different %elution rank thresholds
- Supplementary figure 10 Analysis with MHC I-based binding scores from PRIME2.0: Relative MHC-variant binding vs CH-prevalence
- Supplementary figure 11 Analysis with MHC I-based binding scores from PRIME2.0: Absolute MHC-variant binding vs CH-prevalence
- Supplementary figure 12 Analysis with MHC I-based binding scores from PRIME2.0: Relative MHC-variant binding vs CH clone size
- Supplementary figure 13 Analysis with MHC I-based binding scores from PRIME2.0: Absolute MHC-variant binding vs CH clone size
- Supplementary figure 14 MHC II analysis: Relative MHC-variant binding vs CH-prevalence
- Supplementary figure 15 MHC II analysis: Absolute MHC-variant binding vs CH-prevalence
- Supplementary figure 16 MHC II analysis: Relative MHC-variant binding vs CH clone size
- Supplementary figure 17 MHC II analysis: Absolute MHC-variant binding vs CH clone size
- Supplementary figure 18 Distribution cores for present vs absent variants for each of the 40 variants observed
- Supplementary figure 19 Aggregated scores for present vs absent variants
